# Bile acid metabolites control Th17 and Treg cell differentiation

**DOI:** 10.1101/465344

**Authors:** Saiyu Hang, Donggi Paik, A. Sloan Devlin, Trinath Jamma, Jingping Lu, Soyoung Ha, Brandon N. Nelson, Samantha P. Kelly, Lin Wu, Ye Zheng, Fraydoon Rastinejad, Michael R. Krout, Michael A. Fischbach, Dan R. Littman, Jun R. Huh

## Abstract

Bile acids are abundantly present in the mammalian gut, where they undergo bacteria-mediated transformation, generating a large pool of bioactive molecules. While they have been shown to affect host metabolism, cancer progression and innate immunity, it is unknown whether bile acids affect the function of adaptive immune cells such as T cells expressing IL-17a (Th17 cells) and regulatory T cells (Tregs) that mediate inflammatory and anti-inflammatory responses, respectively. By screening a small-molecule library primarily composed of bile acid metabolites, we identified two distinct derivatives of lithocholic acid (LCA), 3-oxoLCA and isoalloLCA, as specific regulators of Th17 and Treg cells. While 3-oxoLCA inhibited Th17 cell differentiation by directly binding to its key transcription factor RORγt (retinoid-related orphan receptor γ t), isoalloLCA enhanced differentiation of Tregs through mitochondrial-dependent metabolic changes, leading to an increased expression of Foxp3. IsoalloLCA-dependent Treg enhancement required an intronic *Foxp3* enhancer, the conserved noncoding sequence 3 (CNS3), which acts as an epigenetic switch that confers a poised state to the *Foxp3* promoter. Lastly, oral administration of 3-oxoLCA and isoalloLCA to mice led to reduced Th17 and increased Treg cell differentiation in the intestinal lamina propria. Altogether, our data suggest novel mechanisms by which bile acid metabolites control host immune responses by directly modulating the Th17 and Treg balance.

---

Bile acids are cholesterol-derived natural surfactants, produced in the liver and secreted into the duodenum. They are critical for lipid digestion, antibacterial defense and glucose metabolism^1^. While 95% of bile acids are re-absorbed through the terminal ileum of the small intestine and recirculated to the liver, hundreds of milligrams of bile acids are subject to bacterial-mediated transformation and become secondary bile acids with unique chemical structures and biological activities^2,3^. In the healthy human gut, the concentrations of secondary bile acids are in the hundreds of micromolar range^2,4^. While some bile acids disrupt cellular membranes due to their hydrophobic nature^5^, other bile acids protect the gut epithelium^6^ and confer resistance to infection with invasive pathogens such as *Clostridium difficile*^1^. In addition, bile acids were shown to influence gut-associated inflammation, suggesting their potential to regulate gut mucosal immune cells^8,9^. The immune-modulatory effects of bile acids have mostly been studied in the context of innate immunity^10-12^. Although a recent study reported the cytotoxic effects of bile acids on gut-residing T cells^13^, whether they modulate T cell function directly has not been thoroughly examined. Since the identification of digoxin, a plant-derived small molecule harboring a sterol-like core, as the first Th17 cell-specific inhibitor that directly binds to RORγt and inhibits its activity^14^, other structurally similar cholesterol derivatives have been identified as RORγt modulators^15^-^17^. Because bile acids belong to a family of cholesterol metabolites and predominantly exist in the gut where many Th17 cells are present^18^, we reasoned that bile acids might be involved in controlling the function of Th17 cells by modulating RORγt activity.

To identify bile acids with modulatory effects on T cell differentiation, we screened ~30 compounds. We included both primary bile acids, which are synthesized by the host, and secondary bile acids, which are produced by bacterial modification of primary bile acids, in our screen. Naïve CD4^+^ T cells were isolated from wild-type C57BL/6J (B6) mice and cultured with the bile acids under Th17 and, as a counter-screen, Treg differentiation conditions (Extended Data Fig. 1a-c). Strikingly, two distinct derivatives of LCA were found to significantly affect Th17 and Treg cell differentiation. While 3-oxoLCA inhibited Th17 differentiation, isoalloLCA enhanced Tregs, as shown by reduced IL-17a with the former and increased Foxp3 expression with the latter (Fig. 1a, b and Extended Data Fig. 1c). The Treg-enhancing effect of isoalloLCA was particularly evident when T cells were cultured with low levels of TGF-β (Fig. 1a and Extended Data Fig. 2a). Although isoalloLCA strongly enhanced Foxp3 expression in the presence of low, but not high, TGF-β levels (Extended Data Fig. 2a-c), its Treg-enhancing activity requires TGF-β, because pre-treatment of cells with anti-TGF-β antibody prevented Foxp3 enhancement (Extended Data Fig. 2d, e). Modulatory effects of 3-oxoLCA on Th17 cells and isoalloLCA on Tregs were cell-type specific, as neither compound affected T cell differentiation into Th1 or Th2 cells (Fig. 1a, b). While 3-oxoLCA did not affect Treg cell differentiation (Fig. 1b and Extended Data Fig. 1b), isoalloLCA treatment led to ~40-50% reduction of Th17 cell differentiation (Fig. 1a, b). Importantly, both compounds exhibited their T cell modulatory effects in a dose-dependent manner (Fig. 1c). While 3-oxoLCA did not affect cell proliferation, isoalloLCA addition to T cells led to reduced proliferation compared to treatment with DMSO (Extended Data Fig. 2f). IsoalloLCA treatment impaired neither cell viability (Extended Data Fig. 2g) nor T cell receptor (TCR)-mediated activation, as indicated by similar expression levels of TCR activation markers such as CD25, CD69, Nur77 and CD44 (Extended Data Fig. 3a). Treg-enhancement by isoalloLCA required strong TCR activation, as increasing TCR activation with higher concentrations of anti-CD3 resulted in stronger effects on Foxp3 expression without affecting cell viability (Extended Data Fig. 3b-d).

**Figure 1.**
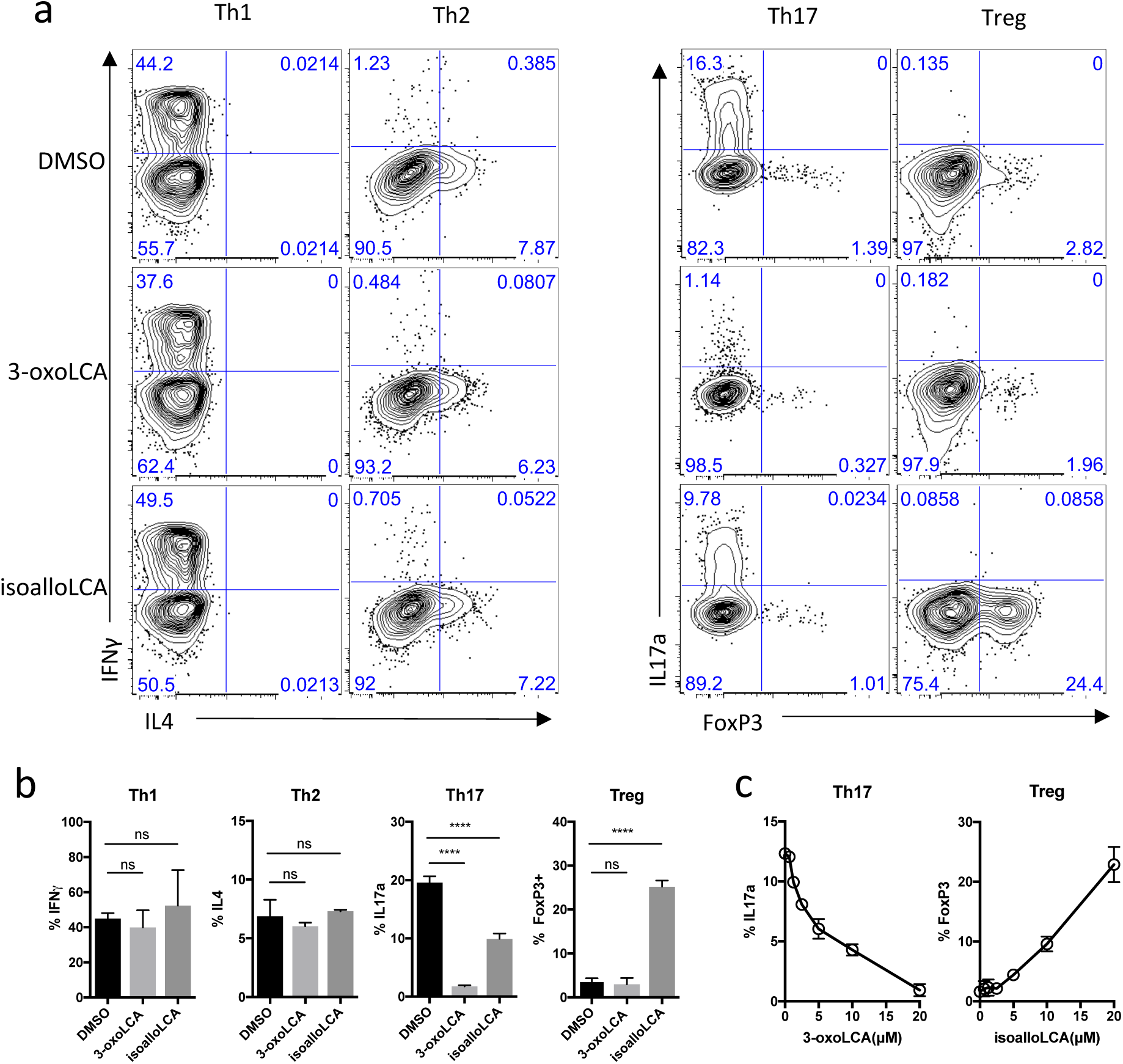
3-oxoLCA inhibits Th17 cell differentiation while isoalloLCA enhances Treg differentiation. **a** and **b**, Flow cytometry and its quantification of intracellular staining for IFN-γ and IL-4 or IL-17a and Foxp3 in sorted naïve T cells activated and expanded in the presence of mouse Th1, Th2, Th17 and Treg polarizing cytokines. A low concentration of TGF-β (0.01 ng/ml) was used for Treg culture. DMSO, 20 μM 3-oxoLCA or 20 μM isoalloLCA was added on day 0 and CD4^+^ T cells were gated for analyses on day 3 for Th17 and Treg, day 5 for Th1 and Th2. c, 3-oxoLCA and isoalloLCA demonstrate dose-dependent effects on Th17 cell and Treg differentiation, respectively. Error bars represent standard deviation. ns; not-significant, ^∗∗∗∗^; p<0.0001, ^∗∗∗^; p<0.001 by unpaired t-test with 2-tailed p-value.

**Figure 2.**
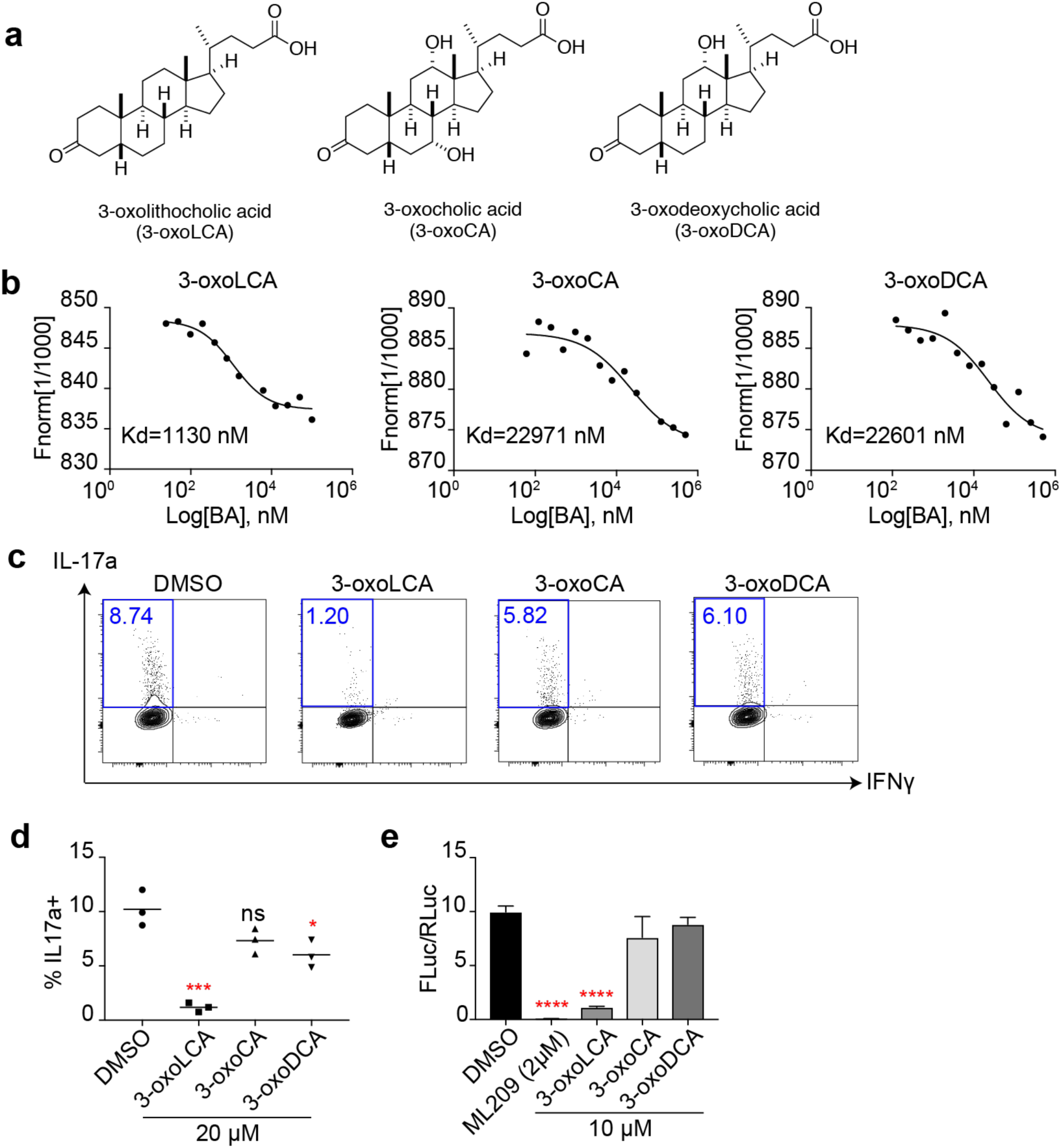
3-oxoLCA physically binds to RORγt and inhibits its transcriptional activities. **a**, Chemical structures of 3-oxoLCA, 3-oxocholic acid (3-oxoCA) and 3-oxodeoxycholic acid (3-oxoDCA). **b**, Microscale thermophoresis assay. 3-oxoLCA binds to RORγ LBD at much lower Kd value than the other two structurally similar bile acids. **c** and **d**, Flow cytometric analyses and quantification of the production of IL-17a from mouse naïve CD4^+^ T cells cultured for 3 days under Th17 polarization condition. DMSO or bile acids at 20 μM were added 18 h after cytokine addition. **e**, 3-oxoLCA, but not 3-oxoCA and 3-oxoDCA, selectively inhibits RORγ-dependent transcriptional activity. Ratio of firefly to Renilla luciferase activity is presented on the y-axis. Error bars represent standard deviation. ns; not-significant, ^∗∗∗∗^; p<0.0001, ^∗∗∗^; p<0.001, ^∗∗^; p<0.01, ^∗^; p<0.01 by upaired t-test with 2-tailed p-value, compared to DMSO control.

To elucidate the mechanisms by which 3-oxoLCA inhibits Th17 cell differentiation, we first examined if 3-oxoLCA physically interacts with the RORγt protein *in vitro*. A microscale thermophoresis (MST) assay was performed with recombinant human RORγt ligand binding domain (LBD). 3-oxoLCA exhibited a robust physical interaction with the RORγt LBD at the equilibrium dissociation constant (Kd) of ~1 μM. We also tested two other structurally similar 3-oxo derivatives of bile acids, 3-oxocholic acid (3-oxoCA) and 3-oxodeoxycholic acid (3-oxoDCA) (Fig. 2a), and demonstrated that these derivatives had ~20 times higher Kd values than 3-oxoLCA (Fig. 2b). In line with this observation, neither 3-oxoCA nor 3-oxoDCA inhibited Th17 cell differentiation as robustly as 3-oxoLCA (Fig. 2c, d). These results suggest that addition of a 3-oxo moiety does not alone confer RORγt-binding activity to bile acids. Next, we examined if 3-oxoLCA can modulate the transcriptional activity of RORγt. As a surrogate reporter for RORγt transcriptional activity, we assayed the effect of the bile acids on firefly luciferase expression directed by a fusion protein of RORγt and Gal4-DBD (DNA binding domain) in human embryonic kidney (HEK) 293 cells. Unlike cells treated with DMSO, cells treated with ML209, a specific RORγt antagonist, completely lost RORγt activity^19^. Likewise, 3-oxoLCA treatment significantly reduced the RORγt reporter activity (Fig. 2e). Altogether, these data suggest that 3-oxoLCA likely inhibits Th17 cell differentiation by physically interacting with RORγt and disabling its transcriptional activity.

We next sought to uncover the mechanism by which isoalloLCA exerts its enhancing effects on Tregs. First, we examined whether other metabolites of LCA can induce Treg cell differentiation. LCA has a 3α-hydroxyl group as well as a cis 5β-hydrogen configuration at the A/B ring junction. This molecule can undergo isomerization, presumably via the actions of gut bacterial enzymes^2^, to form isoLCA (3β,5β), alloLCA (3α,5α) or isoalloLCA (3β,5α) (Fig. 3a). IsoalloLCA, but not the other LCA isomers nor 3-oxoLCA, enhanced Foxp3 expression, confirming that both the 3β-hydroxyl group and *trans* (5α-hydrogen) A/B ring configuration of isoalloLCA are required for Treg enhancement (Fig. 3b). Compared to DMSO-treated cells, isoalloLCA-treated cells significantly inhibited T effector cell proliferation *in vitro*, indicating that they had acquired regulatory activity (Fig. 3c, d). We next investigated whether isoalloLCA enhances Foxp3 expression transcriptionally or post-transcriptionally. T cells isolated from Foxp3-GFP reporter mice exhibited both increased *foxp3* mRNA expression (Fig. 3e) and enhanced GFP levels upon treatment with isoalloLCA (Extended Data Fig. 4a). Thus, isoalloLCA-induced enhanced expression of Foxp3 occurs at the *foxp3* mRNA transcriptional level.

**Figure 3.**
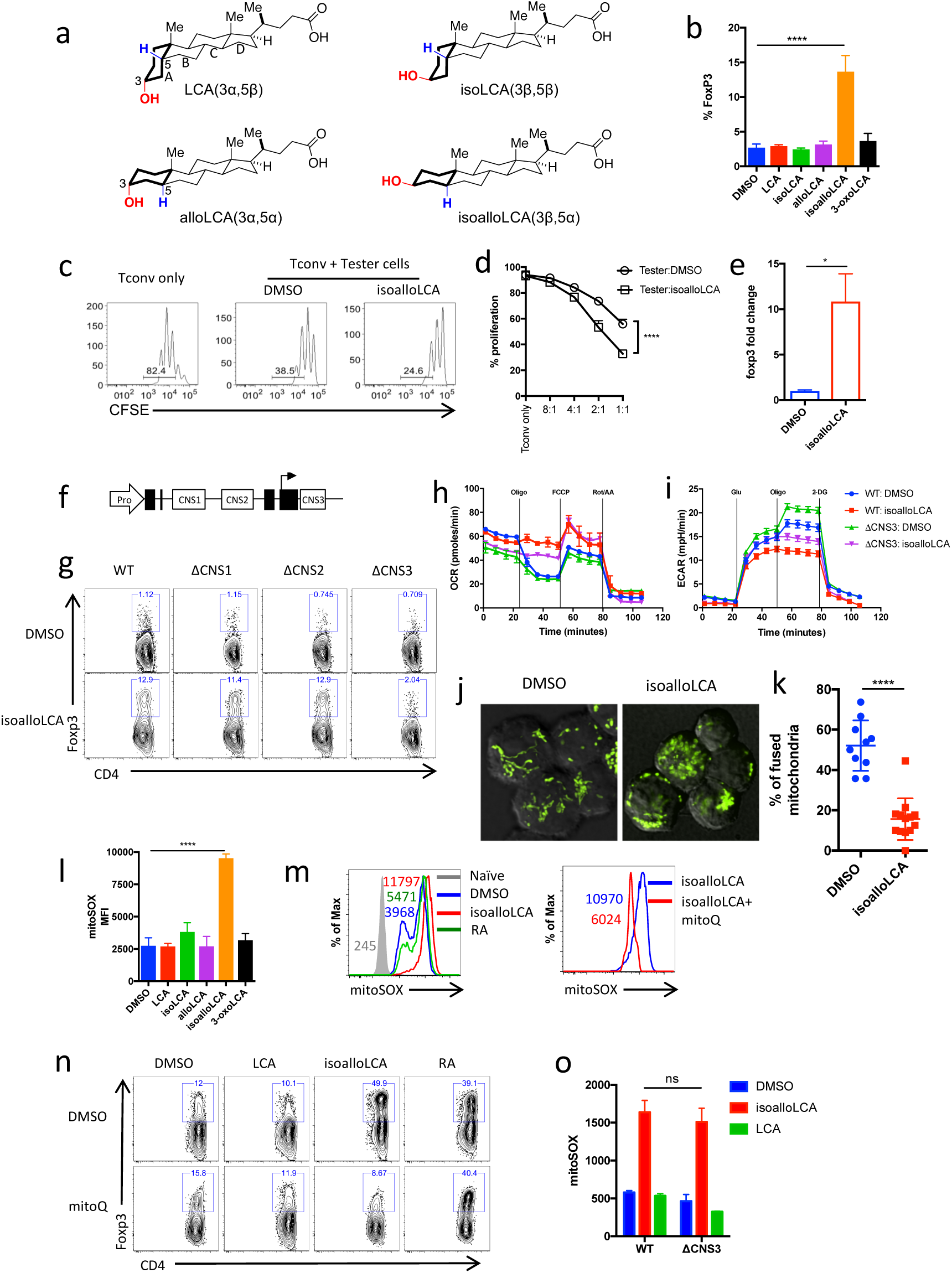
isoalloLCA-dependent enhanced expression of Foxp3 transcription requires both the CNS3 enhancer and increased production of mitoROS. **a**, Chemical structures of LCA isomers: LCA, isoLCA, alloLCA and isoalloLCA. **b**, Foxp3 expression from mouse naïve CD4^+^ T cells cultured for 3 days under Treg polarization conditions with low levels of TGF-β (0.01ng/ml). DMSO or bile acids at 20 μM were added to cell culture. **c** and **d**, *In vitro* suppression assay. CD4^+^ effector T cells were labeled with CFSE and mixed with DMSO- or isoalloLCA-treated Treg cells at different ratios of Tconv: Tester cells. **e**, qPCR analysis for Foxp3 levels in DMSO- or isoalloLCA- (20 μM) treated cells. **f**,Diagram of the Foxp3 gene locus containing the region promoter (Pro) and intronic enhancer regions (CNS1, CNS2 and CNS3). **g**, Flow cytometry of CD4^+^ T cells stained intracellularly for Foxp3. Naïve CD4 T cells isolated from WT, CNS1, CNS2 or CNS3 knockout mice were cultured under Th0 condition (anti-CD3/28 and IL2) in the presence of DMSO and isoalloLCA (20 μM). **h** and **i**, Seahorse analysis of OCR and ECAR with naïve CD4 T cells isolated from WT or CNS3-KO mice cultured under Th0 condition for 48 hours, in the presence of DMSO, isoalloLCA (20 μM). **j** and **k**, Imaging and quantification of mitochondrial morphology of DMSO- or isoalloLCA-treated WT cell. Naïve CD4 T cells from PhaM mice (Jax #018397) were isolated and cultured under Th0 condition (anti-CD3/28 and IL2) for 48 hrs, in the presence of DMSO or isoalloLCA (20 μM). l, Mitochondrial ROS production measured by mitoSOX with T cells, cultured with DMSO or LCA isomers for 48 hrs. MFI denotes mean fluorescence intensity. **m**, Mitochondria ROS production measured by mitoSOX with T cells cultured with DMSO, isoalloLCA (20 μM), retinoic acid (RA), or mitoQ (0.5 μM) for 48 hours. **n**, Representative FACS plots of T cells stained intracellularly for Foxp3, cultured under Treg condition (TGFβ=0.05 ng/ml) in the presence of DMSO, LCA, isoalloLCA (20 μM) or retinoic acid (1 nM), with DMSO or mitoQ (0.5 μM) for 72 hrs. **o**, Mitochondria ROS production measured by mitoSOX with DMSO-, isoalloLCA-, or LCA-treated T cells. Naïve CD4 T cells were isolated from WT or CNS3-KO mice and cultured under Th0 (anti-CD3/28 and IL2) condition. Error bars represent standard deviation. ns; not-significant, ^∗∗∗∗^; p<0.0001, ^∗∗∗^; p<0.001, ^∗∗^; p<0.01, ^∗^; p<0.01 by unpaired t-test with 2-tailed p-value, compared to DMSO control.

The transcriptional regulation of *Foxp3* is governed by three conserved non-coding enhancers, termed CNS1, 2 and 3 (Fig. 3f). These enhancer regions have distinct roles in regulating Treg development, stability and function^220-22^. For example, CNS1 is critical for the generation of peripherally-differentiated Tregs^23^. Treg-promoting small molecules such as the bacterial metabolite butyrate and the vitamin A derivative retinoic acid (RA) enhance Foxp3 expression in a CNSl-dependent manner^24,25^. TGF-β, a key cytokine for Treg differentiation, also partially requires CNS1 for its activity due to the binding of its downstream signaling molecule SMAD3 to the CNS1 enhancer^22,26^. Whereas naïve CD4^+^ T cells from mice with mutations in CNS1 and CNS2 up-regulated Foxp3 as well as wildtype (WT) cells in response to isoalloLCA, cells lacking CNS3 failed to respond (Fig. 3g). This CNS3-dependent effect of isoalloLCA was specific, as other Treg enhancing molecules such as TGF-β and RA boosted Treg differentiation in CNS3-deficient cells, albeit with reduced efficiency (Extended Data Fig. 4b). Thus, unlike other previously identified small molecules that promote Treg cell differentiation in a CNSl-dependent manner, the Foxp3-enhancing activity of isoalloLCA requires CNS3. cRel was previously shown to stimulate Foxp3 expression by directly binding to the CNS3 enhancer^22^. However, both WT and cRel-deficient cells expressed comparable levels of Foxp3 upon isoalloLCA treatment (Extended Data Fig. 4c, d). LCA was shown to activate nuclear hormone receptors, including VDR and FXR^27^. VDR was also implicated in the modulation of both Th17 and Treg cell function^28-31^. Compared to a DMSO-treated control, however, isoalloLCA-treated cells deficient for the expression of VDR or FXR had similar amounts of Foxp3 induction (Extended Data Fig. 5a and c). Thus, CNS3-dependent activation of Foxp3 by isoalloLCA is unlikely to be mediated through the actions of cRel, VDR or FXR. Of note, neither VDR nor FXR contributes to the suppressive activities of 3-oxoLCA on Th17 cells (Extended Data Fig. 5b and d).

CNS3 was previously implicated in Treg cell development by promoting epigenetic modifications such as H3K27 acetylation (H3K27Ac) and H3K4 methylation at the FoxP3 promoter region^21^. Cells treated with isoalloLCA, compared to those treated with DMSO, were found with increased H3K27Ac levels at the FoxP3 promoter region (Extended Data Fig. 6a). Consistent with this, isoalloLCA treatment led to the increased recruitment of histone acetyltransferase p300 to the promoter region (Extended Data Fig. 6b). However, isoalloLCA did not affect H3K4 methylation (Extended Data Fig. 6c). A pan-bromodomain inhibitor iBET that antagonizes H3K27 acetylation prevented, in a dose-dependent manner, isoalloLCA-dependent enhanced expression of Foxp3 (Extended Data Fig. 6d, e). In line with previous work, CNS3 deficiency not only resulted in a significant reduction of overall H3K27Ac levels^21^, but also abrogated the isoalloLCA-dependent increase of histone acetylation level at the FoxP3 promoter region (Extended Data Fig. 6f). Furthermore, p300 recruitment to the Foxp3 promoter region by isoalloLCA was dependent on CNS3 (Extended Data Fig. 6g). Therefore, CNS3 is likely needed to establish a permissible chromatin landscape, whereupon the promoter region can be further acetylated following isoalloLCA treatment through the increased recruitment of p300.

Intricate relationships exist between cellular metabolism and epigenetic modification. For example, substrates for histone acetylation as well as methylation are the byproducts of mitochondrial metabolism^32^. Unlike conventional T cells, Tregs are less dependent on glycolysis but mainly rely on oxidative phosphorylation (OxPhos) for their energy production^33-35^. Indeed, recent studies identified two metabolites, 2-hydroxyglutarate and D-mannose that promote Treg generation by modulating mitochondrial activities^36,37^. To assess whether isoalloLCA affects cellular metabolism such as OxPhos and glycolysis, we measured the oxygen consumption rate (OCR) and extracellular acidification rate (ECAR), respectively, with T cells cultured for 48 h following DMSO or isoalloLCA treatment. At this time point, Foxp3 is not yet strongly induced, thus making it possible to assess isoalloLCA effects on cellular metabolism before cells are fully committed to becoming Tregs. Compared to DMSO, isoalloLCA treatment increased OCR and decreased ECAR, both in WT and CNS3-KO cells (Fig. 3h, i). Notably, isoalloLCA-treated cells were not responsive to oligomycin treatment, which may suggest modified mitochondrial function. Indeed, isoalloLCA-treated cells were found with reduced numbers of fused mitochondria (Fig. 3j, k), which may reflect changes in metabolic activities in mitochondria^38^. Taken together, these findings suggest that isoalloLCA treatment may lead to increased mitochondrial activity. Reactive oxygen species (ROS) are produced as byproducts of mitochondrial OxPhos. Whereas D-mannose was previously shown to increase cytoplasmic ROS production^37^, isoalloLCA treatment led to increased production of mitochondrial ROS (mtROS), without affecting cytoplasmic ROS (Fig. 3l and Extended Data Fig. 6h). Unlike isoalloLCA, other LCA isomers failed to increase mtROS production (Fig. 3l). On the other hand, consistent with the increased mitochondrial respiration, isoalloLCA-treated cells displayed a modest, but significant, increase in mitochondrial membrane potential (Extended Data Fig. 6i). To test if mtROS is directly involved in enhanced Treg differentiation by isoalloLCA, we employed mitoQ, a mitochondrially-targeted antioxidant, to reduce ROS levels in mitochondria (Fig. 3m). Importantly, in the presence of mitoQ, isoalloLCA was no longer effective in enhancing Treg cell differentiation (Fig. 3n). In contrast, RA-dependent induction of Tregs was unaffected by mitoQ treatment (Fig. 3n), thus suggesting distinct modes of action of isoalloLCA and RA. We next investigated if mtROS production is also responsible for the enhanced H3K27Ac levels at the Foxp3 promoter of isoalloLCA-treated cells. Co-treating cells with isoalloLCA and mitoQ led to decreased H3K27Ac levels, compared to those treated with isoalloLCA only (Extended Data Fig. 6j). Therefore, increased mtROS levels likely contribute to isoalloLCA-dependent Treg enhancement. Recently, Foxp3 itself has been shown to enhance mitochondrial OxPhos^39^. In line with this finding, Foxp3-expressing Tregs were found with higher mtROS levels compared to other *in vitro* differentiated CD4^+^ T cell subsets (Extended Data Fig. 6k). Thus, we investigated if increased mtROS production observed in isoalloLCA-treated cells was a secondary effect of enhanced Foxp3 expression. CNS3-deficient cells that did not express high levels of Foxp3 in response to isoalloLCA treatment nevertheless produced increased levels of mtROS as well as enhanced OxPhos (Fig. 3h and o). Taken together, our data thus support a model in which isoalloLCA enhances mtROS production, due to elevated mitochondrial metabolism, which leads to the increased acetylation of H3K27 at the Foxp3 promoter region and enhanced *Foxp3* mRNA transcription (Extended Data Fig. 6l).

We next examined whether 3-oxoLCA and isoalloLCA could influence Th17 and Treg cell differentiation *in vivo* using a mouse model. Segmented filamentous bacteria (SFB), a murine commensal, is known to induce Th17 cell differentiation in the small intestine of B6 mice upon colonization^40^. C57BL/6NTac mice from Taconic Biosciences (Tac) have abundant Th17 cells in their small intestine owing to the presence of SFB. In contrast, C57BL/6J mice from Jackson Laboratories (Jax), which lack SFB, have few intestinal Th17 cells. To first test the *in vivo* effects of 3-oxoLCA in suppressing Th17 cells, we gavaged Jax-B6 mice with an SFB-containing fecal slurry and fed them either a control diet or 0.3% (w/w) 3-oxoLCA-containing chow for a week (Fig. 4a). 3-oxoLCA treatment led to a significant reduction in Th17 cell percentages among total CD4^+^ T cells isolated from the ilea of small intestines (Fig. 4b). SFB colonization levels were comparable between control and 3-oxoLCA-treated groups, suggesting that the change in Th17 cell percentage was not due to a decrease in SFB in the 3-oxoLCA-treated mice (Fig. 4c). In addition, Tac-B6 mice with pre-existing SFB had reduced levels of Th17 cell percentages when fed with 3-oxoLCA compared to control group fed with vehicle. On the other hand, 3-oxoLCA treatment did not affect Treg percentages (Extended Data Fig. 7a-c). Even under gut inflammatory conditions induced by an anti-CD3 injection, which is known to produce robust Th17 cell responses^18,41^, mice treated with 1%, but not with 0.3%, 3-oxoLCA were found with reduced Th17 cell levels (Fig. 4d and Extended Data Fig. 7d). While isoalloLCA alone failed to affect Treg percentages (Extended Data Fig. 7e), mice treated with anti-CD3 antibody and also fed with a mixture of 3-oxoLCA and isoalloLCA were found with increased Treg levels, compared to the vehicle-treated group (Fig. 4e). Altogether, these data suggest that both 3-oxoLCA and isoalloLCA can modulate Th17 and Treg cell responses in mice *in vivo*.

**Figure 4.**
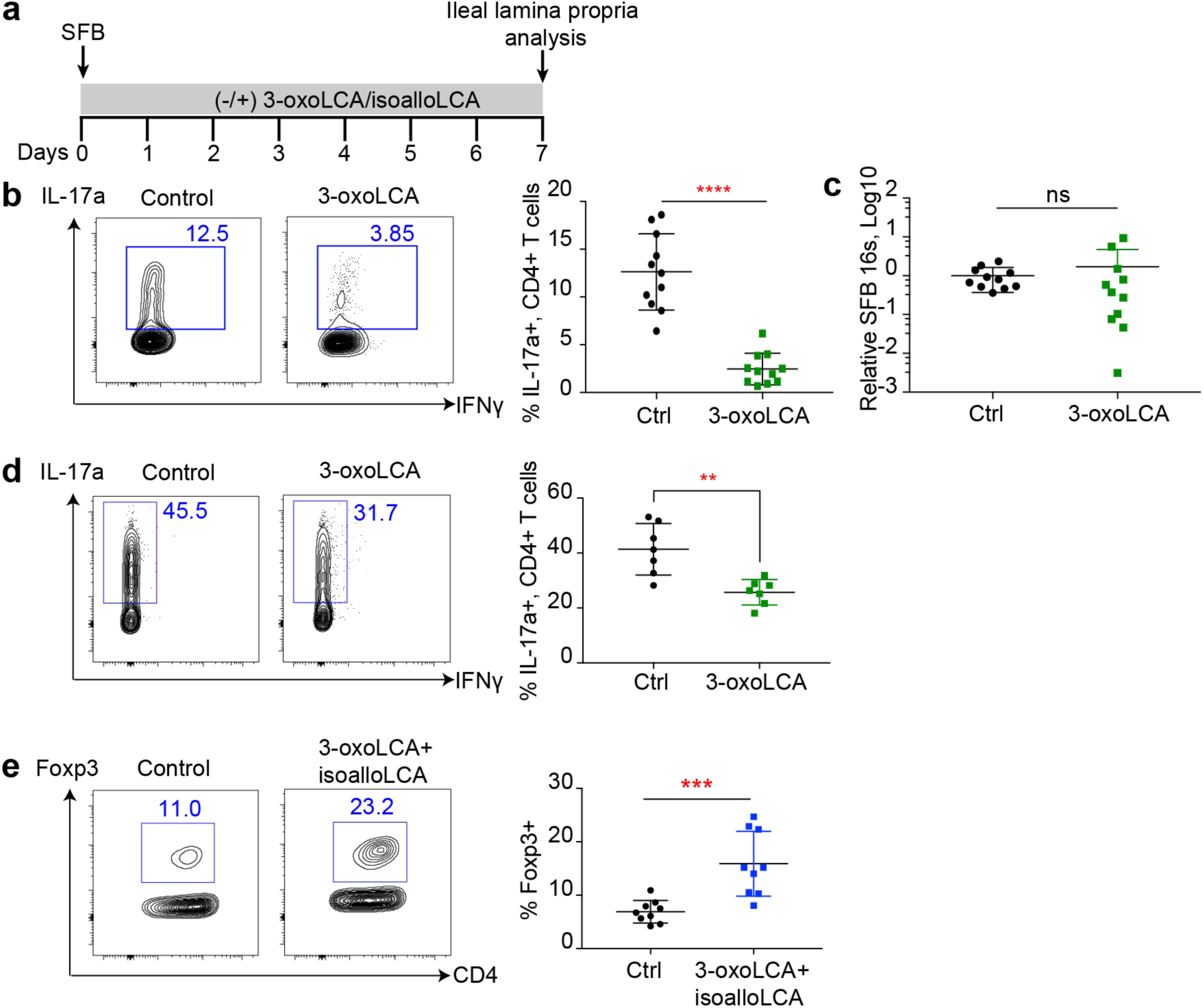
3-oxoLCA inhibits Th17 development while isoalloLCA enhances Treg *in vivo*. **a**, Experimental scheme of Th17 induction by SFB. Jax-B6 animals were fed 3-oxoLCA (0.3%) for a week following acute colonization by SFB. **b**, Flow cytometric analysis of Th17 cells of the small intestinal lamina propria. **c**, SFB colonization measured by qPCR analyses. **d** and **e**, Flow cytometric analysis of Th17 (**d**) and Treg cells (**e**), isolated from ileal lamina propria following anti-CD3 injection, B6 animals were fed 3-oxoLCA (1%) or 3-oxoLCA (1%) + isoalloLCA (0.3%) during the experiments. Error bars represent standard deviation. ns; not-significant, ^∗∗∗∗^; p<0.0001, ^∗∗∗^; p<0.001 by unpaired t-test with 2-tailed p-value.

Lastly, we investigated if *in vitro* treatment of T cells with isoalloLCA produced Tregs exerting suppressive function *in vivo*. The same number of Foxp3+ T cells (CD45.2), sorted from T cell cultures with low or high TGFβ concentrations (TGFβ-lo/-high Tregs) in the absence or presence of isoalloLCA were adoptively transferred into Rag1 KO mice that had also received CD45RB^hi^ naïve CD4^+^ T cells (CD45.1) (Extended Data Fig. 8a, b). Mice that received CD45RB^hi^ or CD45RB^hi^ and TGFβ-lo Tregs developed significant weight loss and shortened colon phenotypes, both of which are indicators of colitis-associated symptoms (Extended Data Fig. 8c, d). On the other hand, adoptive transfer of isoalloLCA-treated, TGFβ-lo Tregs protected mice from developing colitis-associated symptoms to the same degree as those transferred with TGFβ-high Tregs (Extended Data Fig. 8c, d). Furthermore, TGFβ-lo Tregs treated with isoalloLCA were more stable in terms of Foxp3 expression, compared to TGFβ-lo Tregs with DMSO, analyzed eight weeks post transfer (Extended Data Fig. 8e-h). Importantly, mice receiving isoalloLCA-treated TGFβ-lo Tregs had reduced numbers of CD45.1^+^ T effector cells, compared to those with vehicle-treated TGFβ-lo Tregs (Extended Data Fig. 8i). Therefore, isoalloLCA likely promotes stability of Treg cells and enhances their function *in vivo*, leading to decreased proliferation of T effector cells.

Bile acids have been considered as tissue-damaging agents promoting inflammation due to their enhanced accumulation in patients with liver diseases and their chemical properties as detergents that can disrupt cellular membranes^42^. Recent studies, however, have begun to reveal their anti-inflammatory roles, particularly in the innate immune system by suppressing NF-κB-dependent signaling pathways^43,44^ and by inhibiting NLRP3-dependent inflammasome activities^11^. Our studies reveal additional anti-inflammatory roles of two LCA metabolites that directly affect CD4 T cells: 3-oxoLCA suppresses Th17 differentiation while isoalloLCA enhances Treg differentiation. 3-oxoLCA has been reported to exist in both human and rat fecal samples^45,46^, suggesting that gut-residing bacteria may contribute to its production. Given the significant roles of Th17 and Treg cells in a wide variety of inflammatory diseases and their close relationships with gut-residing bacteria, our study suggests the existence of novel modulatory pathways that regulate T cell function and differentiation through microbial metabolites of bile acids. Future studies to elucidate the bacteria and their enzymes that may be responsible for generating 3-oxoLCA and isoalloLCA will provide novel means for controlling Th17 and Treg function in the context of autoimmune diseases and other inflammatory conditions.

## Acknowledgements

We thank Drs. Stephen Smale and Alexander Rudensky for sharing cRel and CNS knockout mice. We also thank Paula Montero Llopis, the Core Director of the MicRoN (Microscopy Resources on the North Quad) at Harvard Medical School, for helping with the mitochondrial imaging. This study was supported by a Charles A. King Trust Fellowship to S.H. and National Institutes of Health grant R01 DK110559 to J.R.H.

## Author Contributions

S.H., D.P., M.A.F., D.R.L., and J.R.H. conceived and designed the experiments; S.H. and D.P. performed most of the experiments; T.J., A.S.D., J.L., S.H., B.N.N., S.P.K., and L.W. provided help with experiments; J.L. and F.R. designed and performed the RORgt binding assay; B.N.N., S.P.K., and M.R.K. synthesized certain bile acid derivatives; Y.Z. provided critical materials; and S.H., D.P., and J.R.H. wrote the manuscript, with contributions from all authors.

## Supplementary Materials

### Materials and Methods

Key Reagents Table.

Key Reagents Table
| REAGENT | SOURCE | IDENTIFIER |
| --- | --- | --- |
| Antibodies |  |  |
| FACS antibody |  |  |
| IFNγ | eBioscience | clone: XMG1.2 |
| IL-4 | Biolegend | clone: 11B11 |
| IL-17a | eBioscience | clone: eBio17B7 |
| Foxp3 | eBioscience | clone: FJK-16s |
| CD4 | eBioscience | clone: RM4-5 |
| CD3e | eBioscience | clone: 145-2C11 |
| CD45 | Biolegend | clone: 30-F11 |
| CD25 | eBioscience | clone: PC61.5 |
| CD69 | eBioscience | clone: H1.2F3 |
| CD44 | Biolegend | clone: IM7 |
| CD62L | eBioscience | clone: MEL-14 |
| CD45RB | Biolegend | clone: C363-16A |
| Nur77 | eBioscience | clone: 12.14 |
| ChIP antibody |  |  |
| Rabbit IgG | abcam | ab46540 |
| H3K27Ac | abcam | ab4729 |
| H3K4me1 | abcam | ab8895 |
| p300 | abcam | ab14984 |
| Cytokines |  |  |
| IL-2 | PEPROTECH | 200-02 |
| IL-4 | R&D systems | 404-ML-010 |
| IL-6 | eBioscience | 14-8061-62 |
| IL-12 | PEPROTECH | 210-12 |
| TGFβ | PEPROTECH | 100-21 |
| hamster IgG | MP Biomedicals | 856984 |
| anti-CD3 | eBioscience | clone: 145-2C11 |
| anti-CD28 | eBioscience | clone: 37.51 |
| anti-IL-4 | eBioscience | clone: 11B11 |
| anti-IFNγ | eBioscience | clone: XMG1.2 |
| anti-TGFβ | eBioscience | clone: 1D11.16.8 |
| IgG1 isotype | eBioscience | clone: P3.6.2.8.1 |

| Chemicals |  |  |
| --- | --- | --- |
| DMSO | Sigma | D8418 |
| 1,25VD3 | Sigma | D1530 |
| Retinoic acid | Sigma | R2625 |
| iBET | Millipore | 401010 |
| Aqua | ThermoFisher | L34957 |
| AnnexinV | eBioscience | BMS306FI/100 |
| CFSE | Biolegend | 423801 |
| mitoQ | Focus Biomolecules | 10-1363 |
| mitoSOX | ThermoFisher | M36008 |
| DCFDA | Sigma | D6883 |
| JC-1 | ThermoFisher | T3168 |
| PMA | Sigma | P1585 |
| Ionomycin | Sigma | I3909 |
| GolgiPlug | BD | 555029 |
| Seahorse drugs |  |  |
| XF mito stress kit | Agilent | 103015-100 |
| XF glycolysis kit | Agilent | 103020-100 |
| Bile Acids |  |  |
| screen library | Michael Fischbach lab |  |
| 3-oxoLCA | Steraloids | C1750-000 |
| isoalloLCA | Steraloids | C0700-000 |
| isoLCA | Steraloids | C1165-000 |
| Bulk chemical synthesis | Michael Krout lab |  |

Mice Table:

Mice Table:
| STRAIN | VENDER | STOCK# |
| --- | --- | --- |
| C57BL/6 | Jackson Laboratory | 000664 |
| C57BL/6 | Taconic Farms |  |
| Foxp3-GFP | Jackson Laboratory | 016958 |
| VDR-KO | Jackson Laboratory | 006133 |
| FXR-KO | Jackson Laboratory | 004144 |
| CD45.1 | Jackson Laboratory | 002014 |
| PhaM<sup>excised</sup> | Jackson Laboratory | 018397 |
| Rag1 KO | Jackson Laboratory | 002216 |
| CNS control | Ye Zheng Lab |  |
| CNS1-KO | Ye Zheng Lab |  |
| CNS2-KO | Ye Zheng Lab |  |
| CNS3-KO | Ye Zheng Lab |  |

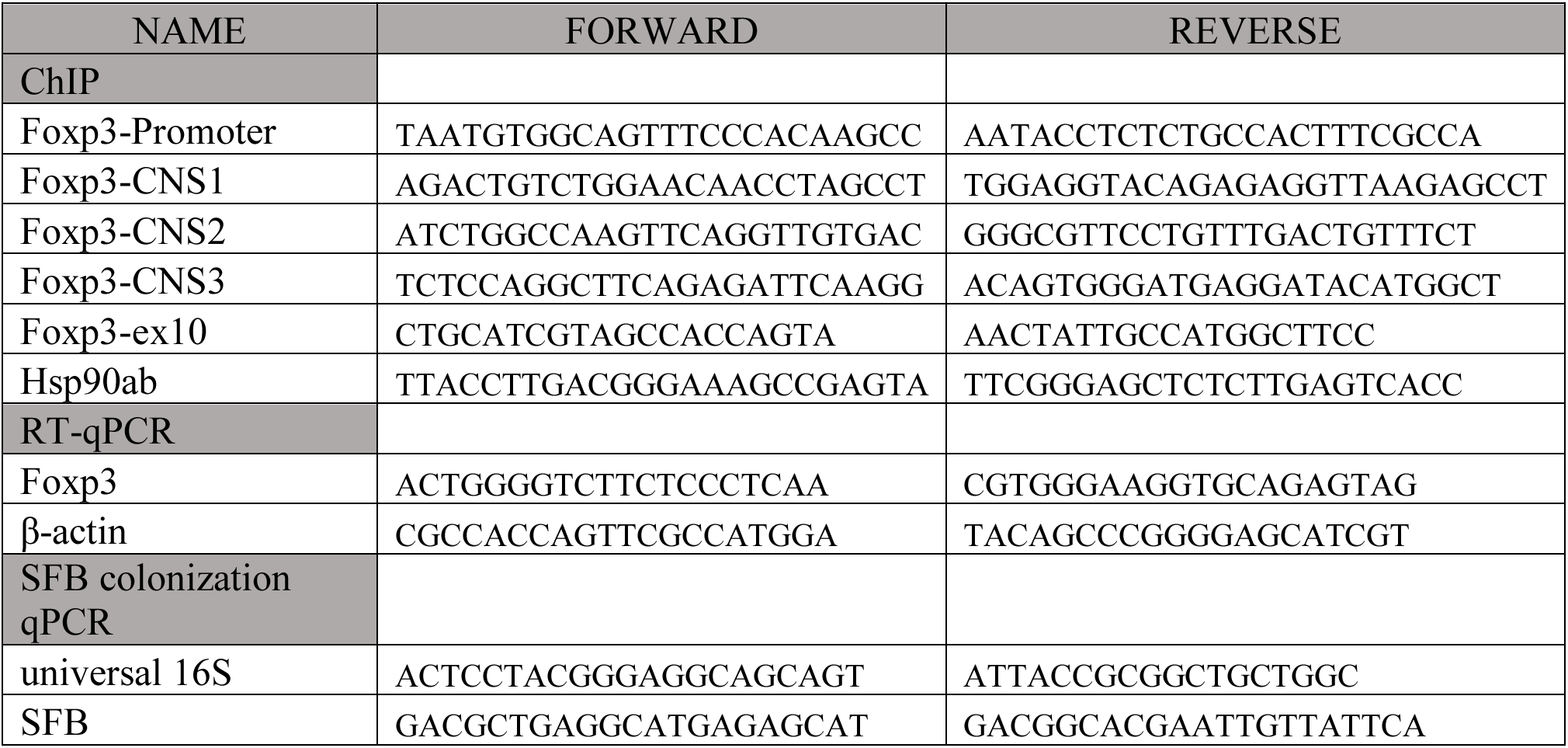
Primers Table:

### Chemical Synthesis of 3-oxoLCA and isoalloLCA

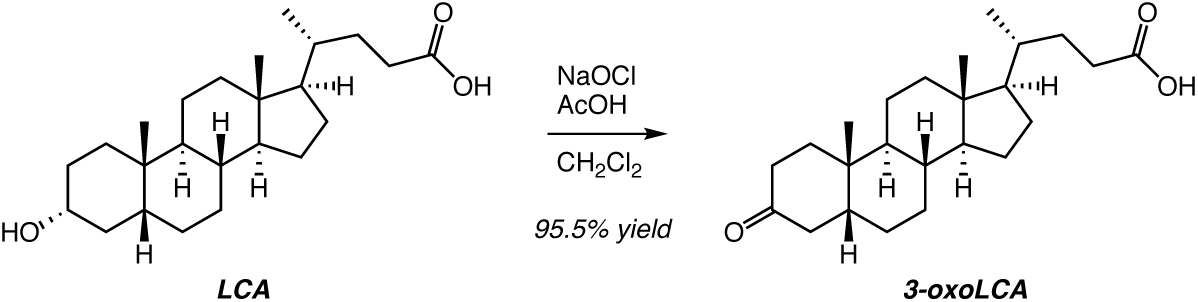

3-oxoLCA was prepared on a large scale by the oxidation of LCA according to the following procedure. Glacial AcOH (160 mL) and CH_2_Cl_2_ (160 mL; total concentration 0.2 M) were added to a 1 L round-bottom flask charged with a stir bar and LCA (24.08 g, 63.95 mmol, 1.0 equivalent). The suspension was stirred until complete dissolution and then immersed in a room temperature water bath. An addition funnel was affixed to the flask and charged with NaOCl (77.6 mL, 95.95 mmol, 1.5 equivalents; 8.25 wt% Chlorox^®^ solution). The bleach solution was added dropwise over a period of ~30 minutes with vigorous stirring. A slight yellowing of the reaction occurred; too rapid of an addition leads to a bright yellow solution and should be avoided. Upon consumption by TLC analysis (~16 h; 19:1 CH_2_Cl_2_/ethyl acetate + 2% AcOH; *p*-Anisaldehyde stain), the excess oxidant was quenched by the addition of *i*-PrOH (24.5 mL, 320 mmol, 5 equivalents). After an additional hour, the reaction was concentrated in vacuo to a slurry. The white slurry was partially dissolved in CH_2_Cl_2_ (400 mL), transferred to a separatory funnel containing a 5% aqueous solution of NaHSO_3_ (150 mL) and acidified to pH < 3 with 1 M HCl. The layers were vigorously mixed, separated, and the aqueous layer was extracted with CH_2_Cl_2_ (3 × 25 mL). The combined organic layers were washed with saturated aqueous NaCl (50 mL), dried over N_2_SO_4_ for >30 minutes, filtered and concentrated in vacuo. The resulting colorless to pale yellow oil forms a white solid over time as the AcOH is removed. The crude ^1^H NMR (CDCl_3_) shows only the desired product with AcOH; however, storage over about a day leads to a noticeable yellowing of the material that increases over time (presumably due to residual oxidant). The crude solid is purified by trituration with Et_2_O/hexanes. To break up any large chunks, Et_2_O (100 mL) and a stir bar are added to the crude solid and stirred. Hexane (100 mL) is then added while stirring, and then the contents were placed in an ice bath and mixing ceased. Vacuum filtration with washing and vacuum drying afforded 3-oxoLCA as a white, crystalline solid (19.488 g, 52.028 mmol, 81.4% yield). The filtrate was concentrated and triturated in a similar manner (30 mL total of 1:1 Et_2_O/hexanes) to generate an additional portion of white crystalline solid (3.3836 g, 9.033 mmol, 14.1% yield). The combined mass of 3-oxoLCA was 22.872 g (61.061 mmol, 95.5% yield).

#### Characterization data

*R_f_* = 0.31 (19:1 CH_2_Cl_2_/ethyl acetate + 2% AcOH; *p*-Anisaldehyde); mp 138–141 °C; ^1^H NMR (600 MHz, CDCl_3_): δ 2.69 (app t, *J* = 14.3 Hz, 1H), 2.40 (ddd, *J* = 15.6, 10.3, 5.2 Hz, 1H), 2.33 (td, *J* = 14.7, 5.3 Hz, 1H), 2.26 (ddd, *J* = 15.8, 9.6, 6.4 Hz, 1H), 2.17 (app d, *J* = 14.2 Hz, 1H), 2.03 (dt, *J* = 8.8, 4.5 Hz, 3H), 1.88 (ddt, *J* = 13.6, 9.2, 4.6 Hz, 2H), 1.83–1.80 (m, 2H), 1.62–1.59 (m, 1H), 1.52–1.06 (m, 15H), 1.02 (s, 3H), 0.94 (d, *J* = 6.5 Hz, 3H), 0.69 (s, 3H); ^13^C{^1^H} NMR (151 MHz, CDCI_3_): δ 213.7, 179.8, 56.6, 56.1, 44.5, 42.9, 42.5, 40.9, 40.2, 37.4, 37.2, 35.7, 35.5, 35.0, 31.1, 30.9, 28.3, 26.8, 25.9, 24.3, 22.8, 21.4, 18.4, 12.2; IR (ATR): 2925, 2878, 2858, 1698, 1376, 1307, 1264, 945, 735 cm^−1^; HRMS (DART–) *m/z:* [M – H]^−^ calculated for C_24_H_37_O_3_ 373.2743, found 373.2752; [α]D^21.5^ +31.5 (*c* 1.205, CH_2_CI_2_). NMR spectral images for 3-oxoLCA are shown in Extended Data Fig. 9 and 10.

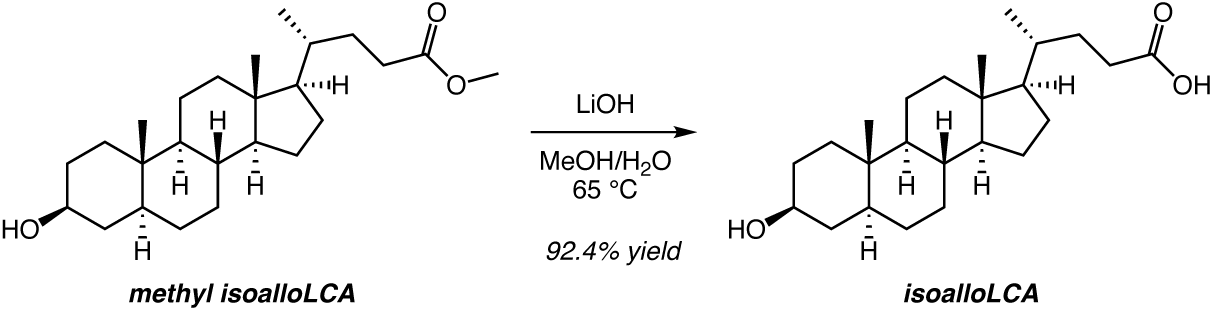

IsoalloLCA was prepared on a large scale by the saponification of methyl isoalloLCA according to the following procedure. Methanol (271 mL) was added to a 1 L round-bottom flask charged with a stir bar and methyl isoalloLCA (6.616 g, 18.894 mmol, 1.0 equivalent). A reflux condenser was affixed to the flask, and the contents were placed under argon and warmed to 65 °C to yield a clear solution. To this solution was added H_2_O (67 mL; 5:1 methanol:H_2_O, 0.05 M) followed by LiOH (2.023 g, 84.47 mmol, 5.0 equivalents) at 65 °C. Upon completion by TLC analysis (2:1 hexanes/ethyl acetate; *p*-Anisaldehyde), the reaction was cooled to room temperature and concentrated in vacuo to a white slurry. H_2_O (200 mL) was added to this slurry, followed by acidification to pH < 3 with 1 M HCl. The congealed mass was sonicated at > 50 °C for at least 3 h to yield a free-flowing, fluffy white solid. The contents were cooled to room temperature, then in an ice bath and isolated via vacuum filtration. Washing and further drying of the filter cake (high vacuum) yielded isoalloLCA as a powdery white solid (5.8797 g, 15.613 mmol, 92.4% yield).

#### Characterization data

*R_f_* = 0.23 (1:1 hexanes/ethyl acetate; *p*-Anisaldehyde); mp 206–210 °C; ^1^H NMR (600 MHz, DMSO-*d*_6_): δ 11.96 (br s, 1H), 4.40 (br s, 1H), 2.22 (ddd, *J* = 15.3, 9.7, 5.4 Hz, 1H), 2.11–2.06 (m, 1H), 1.91 (app d, *J* = 12.3 Hz, 1H), 1.81–1.74 (m, 1H), 1.68–0.80 (m, 24H), 0.86 (d, *J* = 6.6 Hz, 3H), 0.74 (s, 3H), 0.61 (s, 3H), 0.61–0.57 (m, 1H); ^13^C{^1^H} NMR (151 MHz, DMSO-*d*_6_): δ 174.8, 69.3, 56.0, 55.5, 53.8, 44.4, 42.2, 38.2, 36.6, 35.07, 35.04, 34.8, 31.7, 31.3, 30.77, 30.68, 28.4, 27.6, 23.8, 20.8, 18.1, 12.1, 11.9; IR (ATR): 3381, 2922, 2869, 2853, 1700, 1443, 1269, 1191, 1030, 905, 611cm^−1^; HRMS (DART–) *m/z*: [M – H]^−^ calculated for C_2_4H_39_O_3_ 375.2899, found 375.2908; [α]D^22.5^ +28.04 (*c* 0.573, CH3OH). NMR spectral images for isoalloLCA are shown in Extended Data Fig. 11 and 12.

### *In Vitro* T Cell Culture

Naïve CD4^+^ (CD62L^+^ CD44^−^ CD25^−^ CD4^+^) T cells were isolated from the spleens and the lymph nodes of mice of designated genotypes, with FACS sorting. For certain experiments, naïve CD4^+^ T cells were enriched using naïve CD4 T cell isolation kits (Miltenyi). Naïve CD4^+^ T cells (40,000 cells) were cultured in a 96-well plate pre-coated with hamster IgG in T cell medium (RPMI, 10% fetal bovine serum, 25 μM glutamine, 55 μM 2-mercaptoethanol, 100 U/mL penicillin, 100 mg/mL streptomycin) supplemented with 0.25 μg/mL anti-CD3 and 1 μg/mL anti-CD28. For Th0 culture, T cells were cultured with the addition of 100 U/mL of IL-2. For Th1 cell differentiation, T cells were cultured with the addition of 10 μg/mL anti-IL-4 and 10 ng/mL of IL-12. For Th2 cell differentiation, T cells were cultured with the addition of 10 μg/mL of anti-IFNγ and 10 ng/mL of IL-4. For Th17 cell differentiation, T cells were cultured with the addition of 10 ng/mL of IL-6 and 0.5 ng/mL of TGFβ. For Treg culture, T cells were cultured with the addition of 100 U/mL of IL-2 and various concentrations of TGFβ. Bile acids, retinoic acid or mitoQ were added either at 0 or 16 h time points. Cells were harvested and assayed by flow cytometry on day 3. For ROS and mitochondrial membrane potential detection, day 2 cells were incubated with 5 μM of mitoSOX, 10 μM of DCFDA or 2 μM of JC-1 for 30 min and assayed with flow cytometry.

### Flow Cytometry

Cells harvested from *in vitro* culture or *in vivo* mice experiments were stimulated with 50 ng/mL PMA (Phorbol 12-myristate 13-acetate) and 1 μM ionomycin in the presence of GolgiPlug for 4hr to determine cytokine expression. After stimulation, cells were stained with cell surface marker antibodies and LIVE/DEAD Fixable dye, Aqua, to exclude dead cells, fixed and permeabilized with a Foxp3/Transcription factor staining kit (eBioscience, #00552300), followed by staining with cytokine-specific antibodies. All flow cytometry analyses were performed on an LSR II flow cytometer (BD) and data were analyzed with FlowJo software (TreeStar).

### Cell Proliferation Assay

Naïve CD4^+^ T cells were labeled with 1 μM carboxyfluorescein succinimidyl ester (CFSE) and cultured for 3 days prior to FACS analysis.

### *In Vitro* Suppression Assay

A total of 2.5×10^4^ freshly-purified naïve CD4^+^CD25^−^CD44^−^CD62L^hlgh^ T cells (Tconv) from CD45.1 B6 mice were labeled with 1 μM CFSE, activated with soluble anti-CD3 (1 μg/mL) and 5×10^4^ APCs in 96-well round-bottom plates for 3 days in the presence of tester cells (CD45.2). The CFSE dilution of CD45.1 Tconv cells was assessed by flow cytometry.

### Mammalian Luciferase Reporter Assay

Reporter assays were conducted as previously described ^1^. Briefly, 50,000 human embryonic kidney 293T cells per well were plated in 96-well plates in antibiotic-free Dulbecco’s Modified Eagle Media (DMEM) containing 1% fetal calf serum (FCS). Cells were transfected with a DNA mixture containing 0.5 μg/mL of firefly luciferase reporter plasmid (Promega pGL4.31), 2.5 ng/mL of a plasmid containing Renilla luciferase (Promega pRL-CMV), and G4DBD-RORγ (0.2 μg/mL). Transfections were performed using TransIT-293 (Mirus) according to the manufacturer’s instruction. Bile acids or vehicle control were added 24 h after transfection and luciferase activity was measured 16 h later using the dual luciferase reporter kit (Promega).

### Microscale Thermophoresis Assay

The binding affinity of the compounds with RORγ-LBD was analyzed by Microscale thermophoresis (MST). Purified RORg-LBD was labeled with the Monolith NT^™^ Protein Labeling Kit RED (NanoTemper Technologies). Serially-diluted compounds, with concentrations of 1 mM to 20 nM, were mixed with 55 nM labeled RORγ-LBD at room temperature and loaded into Monolith TM standard-treated capillaries. Binding was measured by monitoring the thermophoresis with 20% LED power and ‘Medium’ MST power on a Monolith NT.115 instrument (Nano Temper Technologies) with the following time setting: 5s Fluo, Before; 20s MST On; 5s Fluo, After. Kd values were fitted using the NT Analysis software (Nano Temper Technologies).

### RT-qPCR

Total RNA was isolated from cultured T cells using an RNeasy kit (Qiagen, #74134) and reverse transcribed using a PrimeScript RT kit (Takara, #RR037A). Primers used for qPCR are listed in the primer table. All qPCRs were run on the Bio-Rad CFX real-time system using iTaq Universal SYBR Green Supermix (Bio-Rad, #1725124). β-actin was used as an internal control to normalize the data across different samples.

### Metabolic Assays

*In vitro* differentiated cells were cultured in the presence of DMSO or isoalloLCA for 48 h. Oxygen consumption rate (OCR) and extracellular acidification rate (ECAR) were determined using a Seahorse XF96 Extracellular Flux Analyzer (Seahorse Bioscience) following protocols recommended by the manufacturer and according to the previously published method ^2^. Briefly, cells were seeded on XF96 microplates (150,000 cells/well) that had been pre-coated with poly-D-lysine (Sigma) to immobilize cells. Cells were maintained in XF medium containing DMSO or isoalloLCA in a non-CO2 incubator for 30 min before the assay. ECAR was measured with an XF glycolysis stress test kit with sequential injection of 10 mM glucose, 2 μM oligomycin and 50 mM 2-DG (2-deoxy-D-glucose). The Mito stress test kit was used to test OCR by sequential injection of 1 μM oligomycin, 1.5 μM FCCP and 0.5 μM rotenone/antimycin A. Data were analyzed by wave software (Agilent).

### Mitochondrial Imaging and Quantification

Naïve CD4^+^ T cells derived from PhaM^excised^ mice ^3^ were cultured in the presence of DMSO or isoalloLCA for 2 days. Cells were then harvested and fixed with 4% paraformaldehyde for 20 mins, washed, immobilized on 2% agarose in PBS pads and covered with #1.5 coverslips. Fluorescence images were acquired using a motorized Nikon Ti inverted microscope equipped with a Yokogawa CSU-W1 spinning disk confocal head with a 50 *μ*m pinhole size, an Andor Zyla 4.2 plus sCMOS camera, Toptica iChrome MLE 4-color multi-laser engine (488, 515, 561 and 640 nm), SOLA 395 engine widefield illuminator, and Nikon motorized stage with a PI 250 μm piezo insert, using a Plan Apo 100x/1.45 DIC objective. Fluorescence imaging was performed using a Chroma quad multipass dichroic mirror and Chroma 525/36 nm bandpass emission filter. The acquisition software was NIS Elements AR 5.02.

### Chromatin immunoprecipitations

Chromatin immunoprecipitation (ChIP) assays were performed according to standard protocol. Briefly, naïve CD4^+^ T cells were cultured for 48 h, and fixed for 10 min with 1% formaldehyde. Then 0.125 M glycine was added to quench the formaldehyde. Cells were lysed, and chromatin was harvested and fragmented by sonication at a concentration of 10^7^ cells/ChIP sample. Chromatin was immunoprecipitated with 5 μg of ChIP or IgG control antibodies at 4 °C overnight and incubated with protein G magnetic beads (ThermoFisher, # 10001D) at 4 °C for 2 h, washed, and eluted in 150 μL elution buffer. Eluate DNA and input DNA were incubated at 65 °C to reverse the crosslinking. After digestion with proteinase K, DNA was purified with the QIAquick PCR purification kit (Qiagen, # 28104). The relative abundance of precipitated DNA fragments was analyzed by qPCR using SYBR Green Supermix (Bio-Rad, #1725124). Primer information is listed in the table.

### Isolation of lamina propria lymphocytes

Gut tissues were harvested and treated with 1 mM DTT at room temperature for 10 min, and 5 mM EDTA at 37 °C for 20 min to remove epithelial cells, and dissociated in digestion buffer (RPMI, 1 mg/mL collagenase, 100 μg/mL DNase I, 5% FBS) with constant stirring at 37 °C for 30 min. Mononuclear cells were collected at the interface of a 40%/80% Percoll gradient (GE Healthcare). Cells were then analyzed by flow cytometry.

### Animal experiments

For bile acid feeding experiments, the standard mouse diet in ground meal format (Picolab Diet, #5053) was evenly mixed with a measured amount of bile acid compounds and provided in glass feeder jars and replenished when necessary. Colonization of mice with SFB was done with fresh fecal samples, derived from *il23r*; *rag2* double-knockout mice that are known to carry much higher levels of SFB compared to conventional B6 mice. Fecal samples were homogenized in water using a 70 μm cell strainer and a 5 mL syringe plunger. Supernatant was introduced into mice using a 20G gavage needle at 250 μL/animal, approximately equal to the amount of 1/4 mouse fecal pellets.

### Adoptive Transfer Colitis

CD45RB^hi^ adoptive transfer colitis was performed as described ^4^ Briefly, isolated CD4^+^CD25^−^CD45RB^hi^ naïve T cells were sorted from wild-type B6.CD45.1 mice by FACS and 0.5 million cells were adoptively transferred into each Rag1-KO recipient mouse. The same number of *in vitro* cultured cells (CD45.2) were co-transferred to the recipient mice. Mice were then monitored and weighed each week. At week 8, colon tissues were harvested, and lamina propria lymphocytes were analyzed by flow cytometry.

### Statistical

Statistical analysis tests were performed with Prism (GraphPad).

---

**Extended Data Figure 1.**
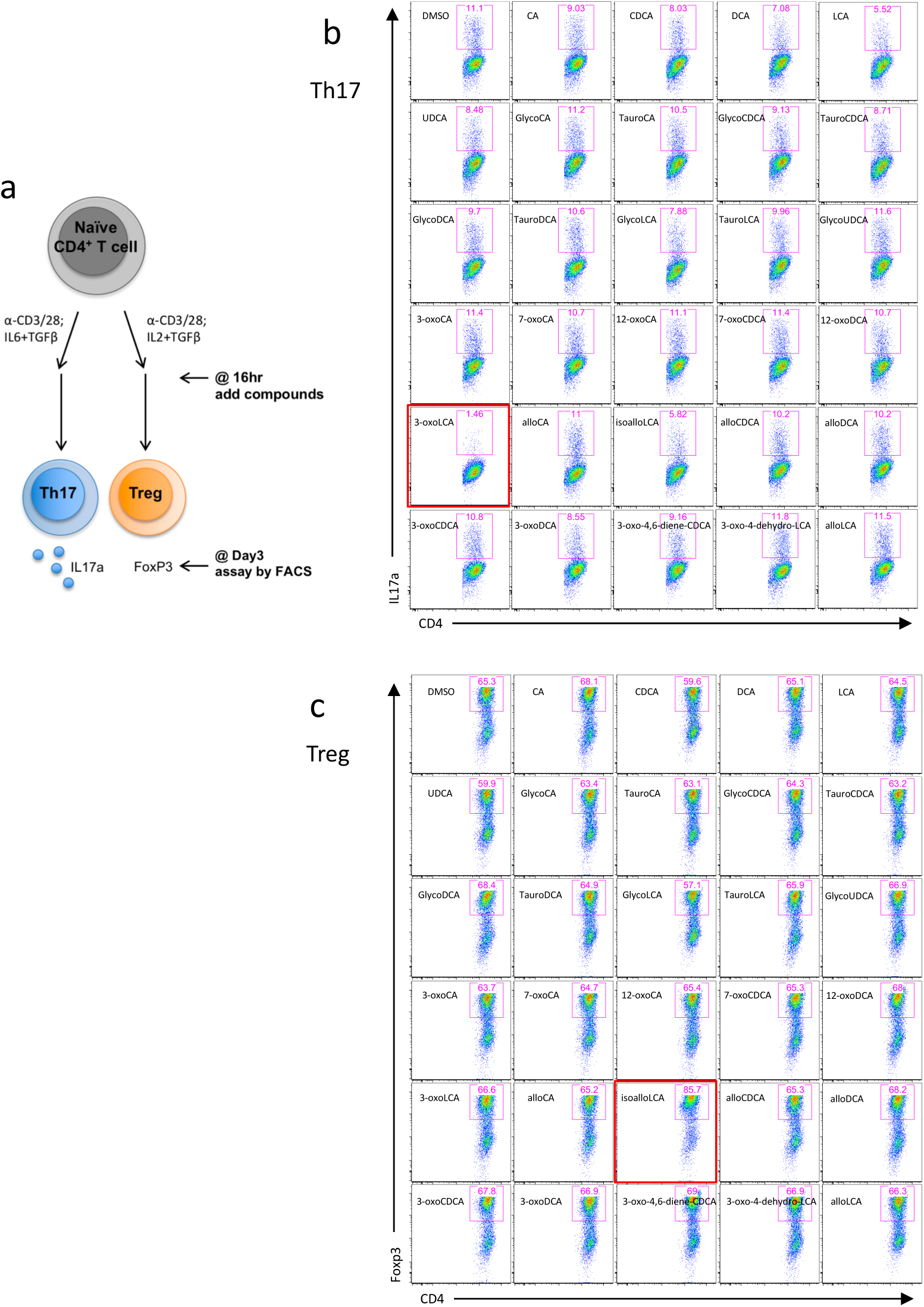
3-oxoLCA and isoalloLCA affect Th17 and Treg differentiation. **a**, Schematic of the screening procedure, **b** and **c**, Naïve CD4^+^ T cells isolated from B6 mice were cultured under Th17 (IL6=10 ng/ml; TGFβ=0.5 ng/ml) (**b**) and Treg (IL2=100 U/ml; TGFβ=0.1 ng/ml) (**c**) polarization conditions for 3 days. DMSO or various bile acids at 20 μM concentration were added to the cell cultures on day 1. Representative FACS plots of CD4^+^ T cells stained intracellularly for IL-17a and Foxp3.

**Extended Data Figure 2.**
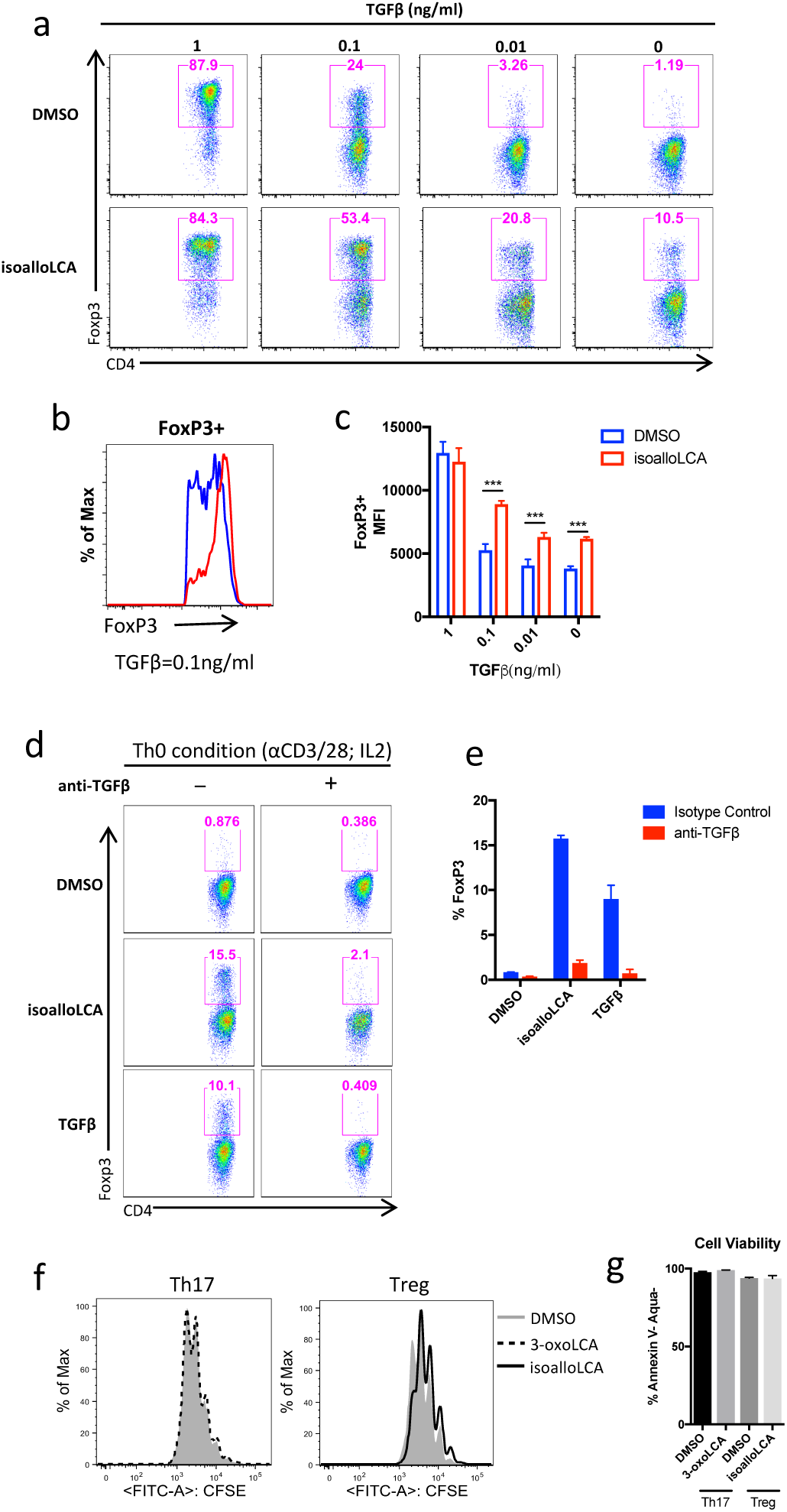
isoalloLCA-induced Treg expansion requires TGF-β. **a-c**, Flow cytometry and histogram of CD4^+^ T cells, cultured for 3 days with different amounts of TGF-β (1; 0.1; 0.01; 0 ng/ml) and IL-2 (100 U/ml) in the presence of DMSO or isoalloLCA (20 μM) and intracellularly stained for Foxp3. MFI denotes mean fluorescence intensity (n=3 per group). ^∗∗∗^p<0.001 calculated by one-way ANOVA. **d** and **e**, Flow cytometry of CD4^+^ T cells, cultured for 3 days in the presence of DMSO, isoalloLCA (20 μM) or TGF-β (0.05 ng/ml). In addition, anti-TGFβ antibody (10 μg/ml, 1D11) or isotype control was added to the culture, **f**, Naïve CD4^+^ T cells were labeled with a cell proliferation dye CFSE and cultured for 3 days in the presence of DMSO, 3-oxoLCA or isoalloLCA under Th17 or Treg polarization condition, **g**, Live cell percentages at the end of the 3-day culture were determined based on both Annexin V and Aqua staining.

**Extended Data Figure 3.**
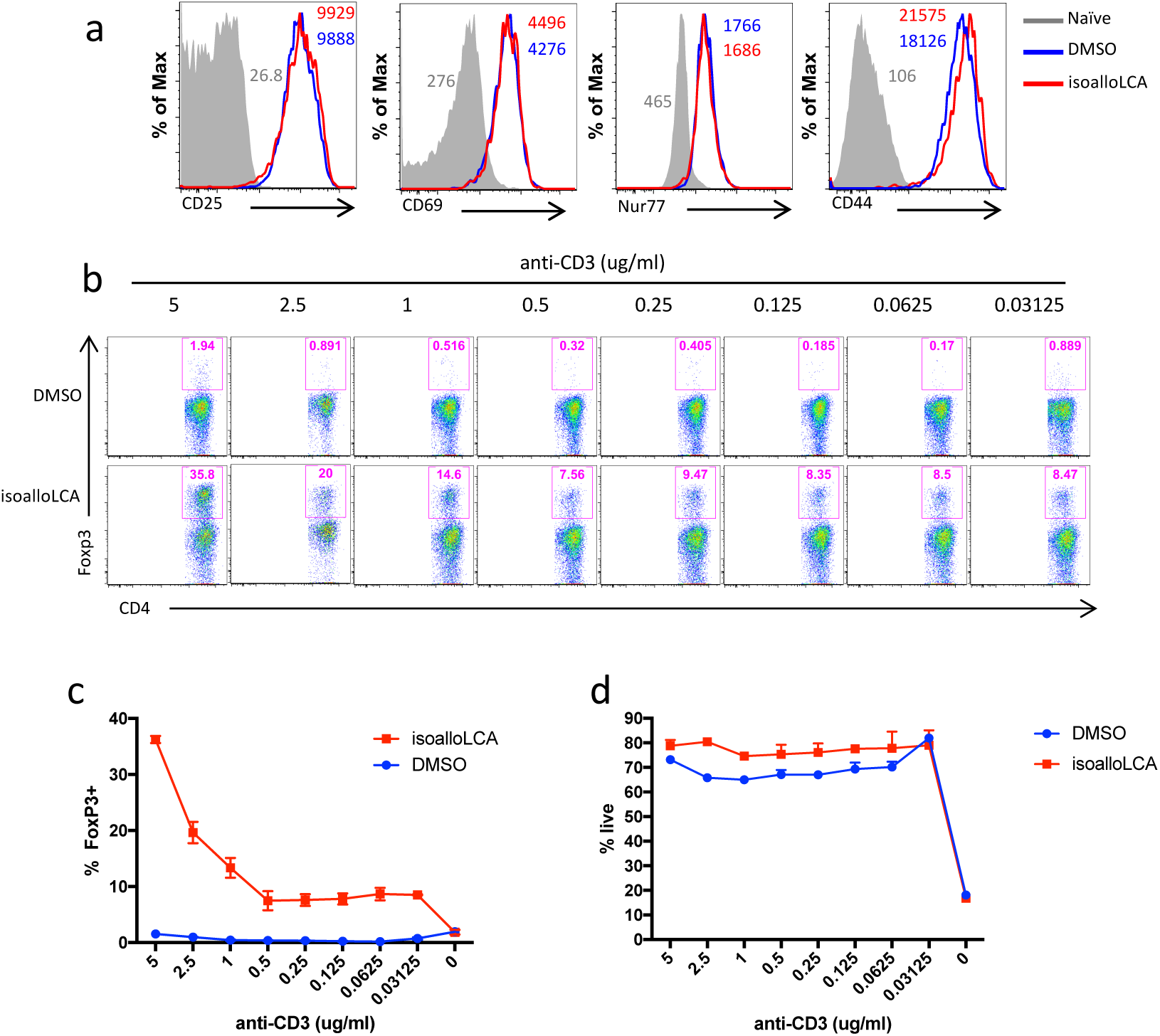
isoalloLCA does not affect T cell activation but its effects on Foxp3 expression require strong TCR stimulation. **a**, Both DMSO and isoalloLCA treatment lead to comparable levels of expression of CD25, CD69, Nur77 and CD44. Naïve CD4^+^ T cells were used as a negative control, **b-d**, T cells were cultured under Th0 condition with different concentrations of anti-CD3ε antibody, in the presence of DMSO or isoalloLCA (20 μM). Representative FACS plots of CD4^+^ T cells cultured for 3 days and stained intracellularly for Foxp3 (**b**). Quantification of Foxp3^+^ and viable T cells after 3-day culture (**c** and **d**).

**Extended Data Figure 4.**
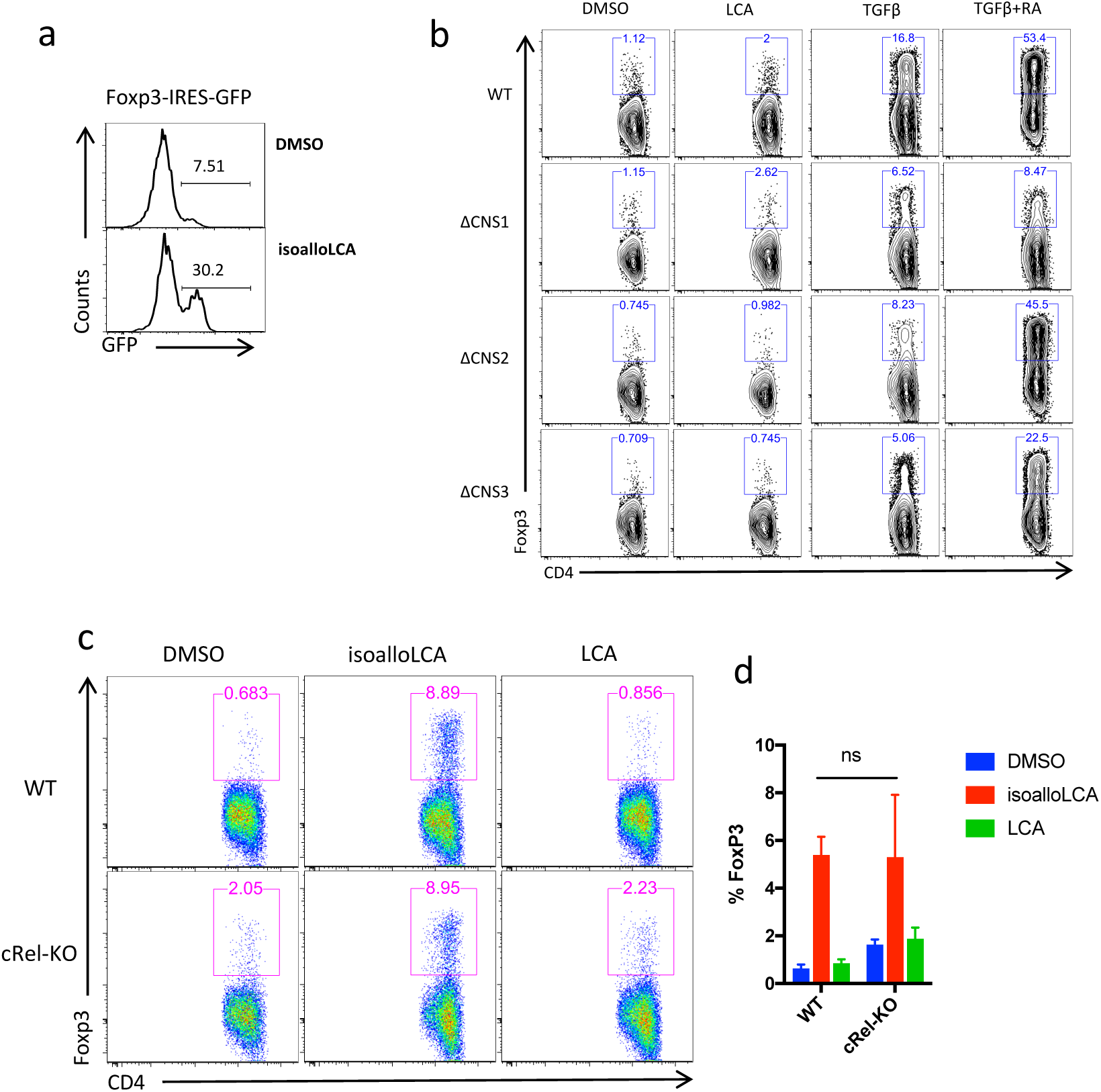
cRel is dispensable for isoalloLCA-dependent induction of FoxP3. **a**, Expression of GFP in DMSO- or isoalloLCA-treated T cells cultured under Treg condition with low concentration of TGFβ (0.01 ng/ml). Naïve CD4^+^ T cells were isolated from FoxP3-IRES-GFP mice. **b**, Flow cytometry of CD4^+^ T cells stained intracellularly for Foxp3. Naïve CD4 T cells isolated from WT, CNS1, CNS2 or CNS3 knockout mice were cultured under Th0 condition (anti-CD3/28 and IL2) in the presence of TGFβ (0.05 ng/ml) and additional retinoic acid (RA; 1 ng/ml). **c** and **d**, Flow cytometry **c**) and its quantification (**d**) of CD4^+^ T cells stained intracellularly for Foxp3. Naïve CD4 T cells were isolated from WT control mice or cRel-KO mice and cultured under Th0 condition in the presence of DMSO, isoalloLCA (20 μM) or LCA (20 μM). Error bars represent standard deviation. ns, not-significant, by unpaired t-test with two-tailed p-value.

**Extended Data Figure 5.**
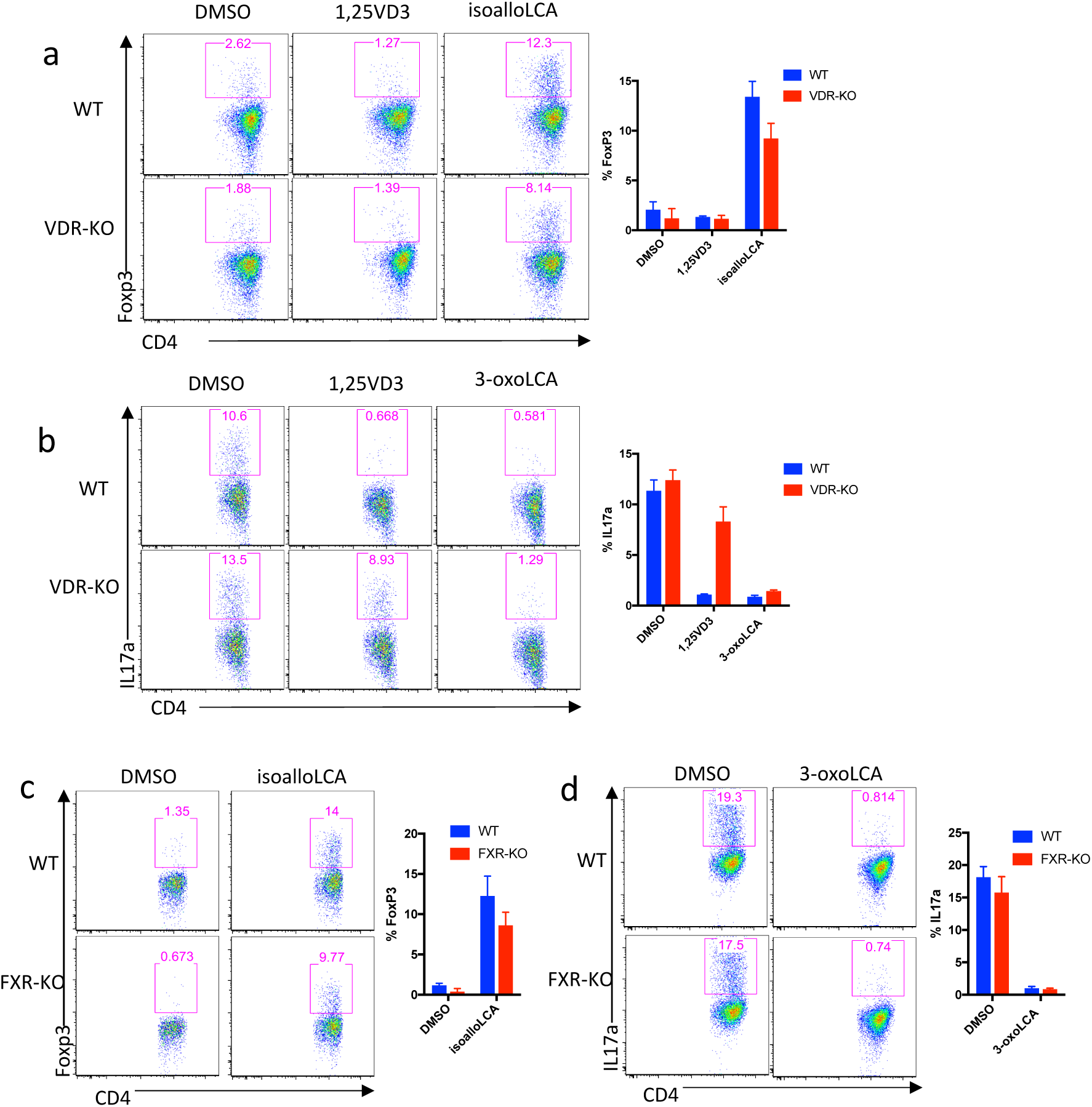
VDR and FXR are not the targets of 3-oxoLCA and isoalloLCA. **a** and **b**, Naïve CD4 T cells isolated from WT control or VDR knockout mice were cultured under Treg (**a**) or Th17 (**b**) conditions for 3 days, in the presence of DMSO, VDR agonist 1,25-dihydroxyvitamin D3 (1,25VD3, 10nM), 3-oxoLCA (20 μM) or isoalloLCA (20 μM). Representative FACS plots of T cells intracellularly stained for Foxp3 or IL17a. **c** and **d**, Naïve CD4 T cells isolated from WT control or FXR knockout mice were cultured under Treg (**c**) or Th17 (**d**) conditions for 3 days, in the presence of DMSO, 3-oxoLCA (20 μM) or isoalloLCA (20 μM). Representative FACS plots of T cells intracellularly stained for Foxp3 or IL17a.

**Extended Data Figure 6.**
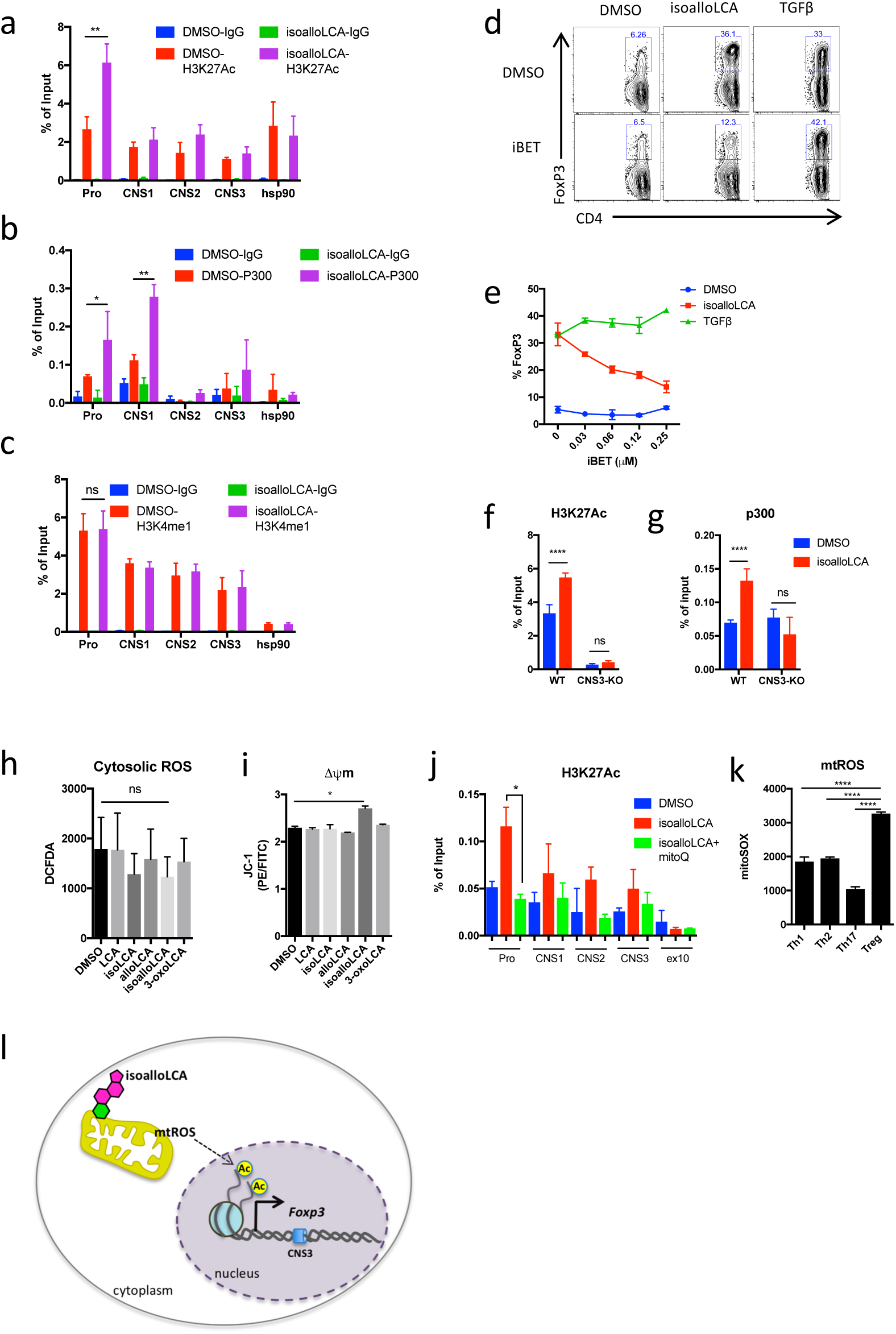
IsoalloLCA dependent Foxp3 transcription requires mitochondrial ROS and H3K27Ac. **a-c**, ChIP analysis of H3K27Ac, p300 and H3K4me1 on Foxp3 gene locus. Chromatin obtained from DMSO- and isoalloLCA treated WT cells was immunoprecipitated with IgG, anti-H3K27Ac, anti-P300, or anti-H3K4me1 antibodies, followed by real-time PCR analysis. Primers targeting foxp3 promoter (Pro), CNS1, CNS2 and CNS3 region and hsp90 promoter were used for qPCR quantification. **d** and **e**, Flow cytometry and quantification of CD4^+^ T cells stained intracellularly for Foxp3. Naïve CD4 T cells isolated from WT mice were cultured under Th0 condition (anti-CD3/28 and IL2) in the presence of DMSO (20 μM), isoalloLCA (20 μM) or TGFβ (0.1 ng/ml) in the presence or absence of iBET (0.25 μM). **f** and **g**, ChIP analysis of H3K27Ac and p300 on the Foxp3 promoter region. Naïve CD4^+^ T cells isolated from WT or CNS3 knockout mice were treated with DMSO or isoalloLCA. **h** and **i**, Cytoplasmic ROS measured by 2’,7’-dichlorofluorescin diacetate (DCFDA) and mitochondrial membrane potential measured by JC-1 dye with T cells cultured with DMSO or LCA, isoLCA, alloLCA, isoalloLCA or 3-oxoLCA at 20 μM for 48 hours. **j**, ChIP analysis of H3K27Ac on the Foxp3 promoter of T cells, treated with DMSO, isoalloLCA or isoalloLCA+mitoQ for 72 hrs. **k**, Mitochondria ROS production measured by mitoSOX with T cells cultured under Th1, Th2, Th17 or Treg condition for 3 days. **l**, A model showing the mechanism of isoalloLCA enhancement of Treg differentiation. Error bars represent standard deviation. ns; not-significant, ^∗∗∗∗^; p<0.0001, ^∗∗∗^; p<0.001 by unpaired t-test with two-tailed p-value.

**Extended Data Figure 7.**
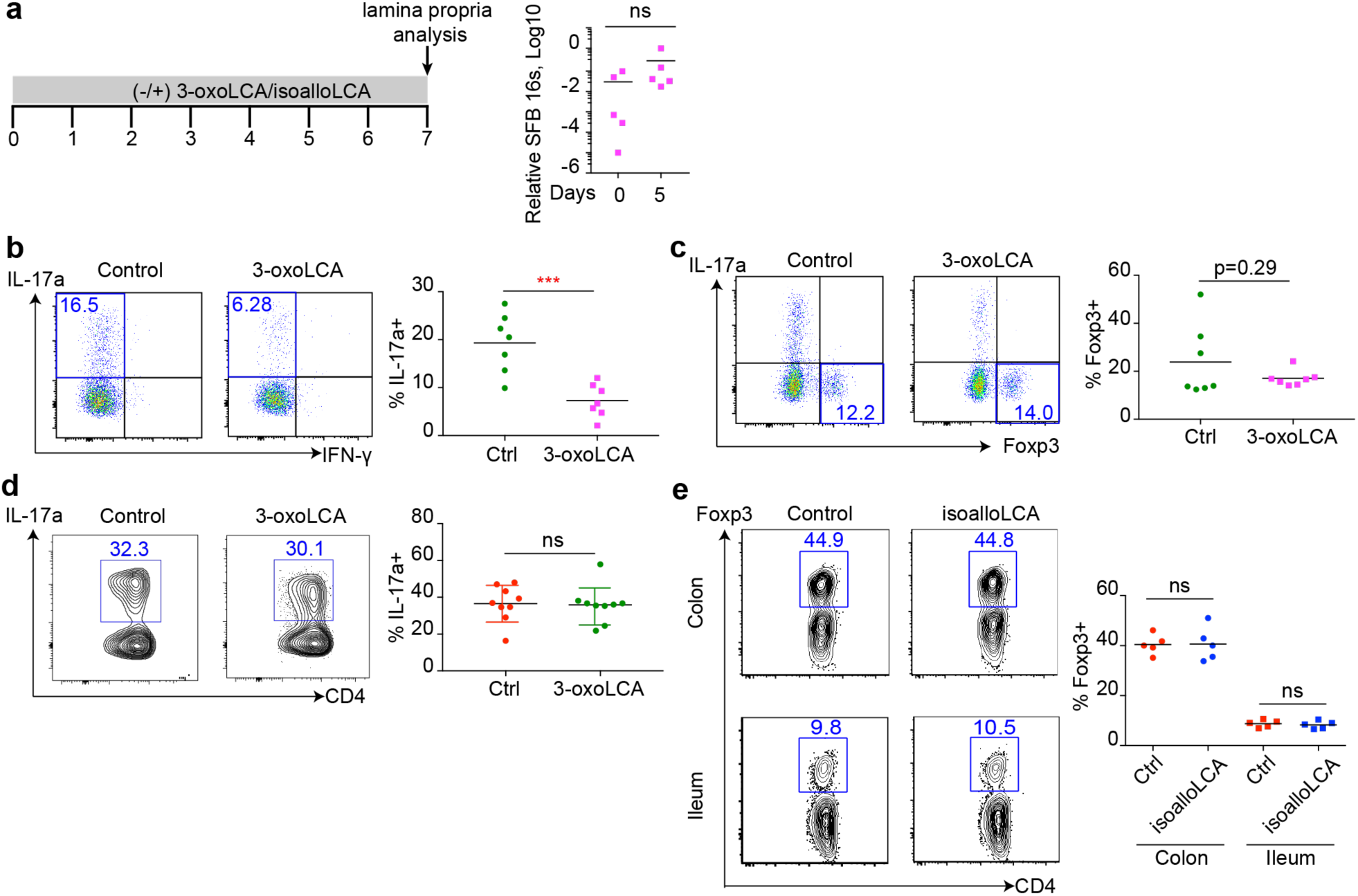
3-oxoLCA inhibits Th17 cell in steady state. **a**, Experimental scheme. SFB colonization measured by qPCR analysis. **b** and **c**, Flow cytometric analysis and quantification of CD4^+^ cells of the small intestinal lamina propria from B6 mice with pre-existing SFB treated with vehicle or 3-oxoLCA (0.3%) in the diet. **d** and **e**, Flow cytometric analysis and quantification of CD4^+^ cells of the lamina propria following anti-CD3 injection, B6 mice were treated with vehicle or isoalloLCA (0.3%) in the diet. ^∗∗∗^p<0.001; ns, not significant by unpaired t-test with two-tailed p-value.

**Extended Data Figure 8.**
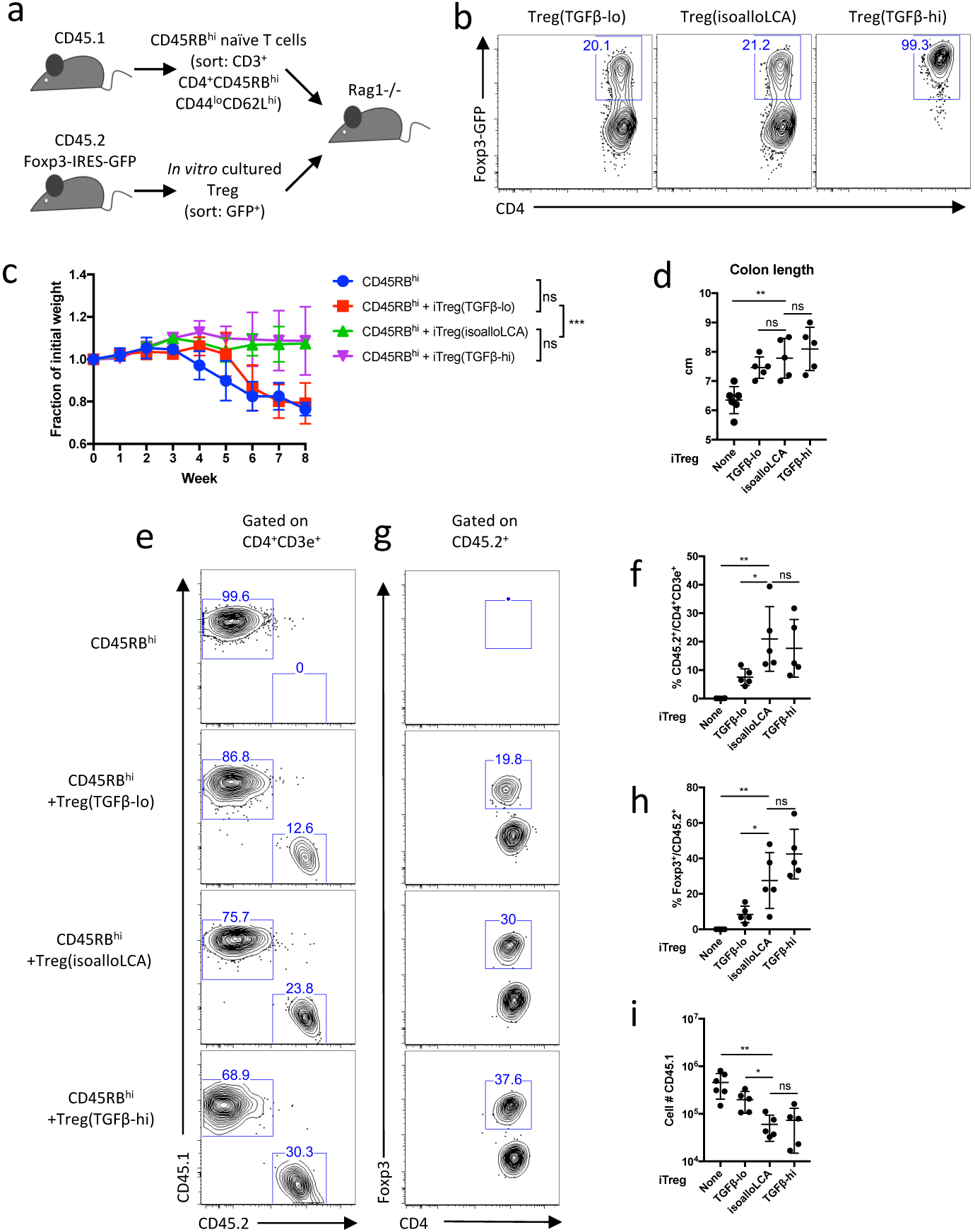
isoalloLCA induced Treg cells suppress transfer colitis. **a**, Experimental scheme. Rag1^-/-^ recipient mice were transferred intraperitoneally with 0.5 million CD45RB^hi^ naïve CD4 T cells from CD45.1 mice, with or without co-transfer of 0.5 million Foxp3-GFP^+^ Treg cells. Foxp3-GFP^+^ cells were cultured under TGFβ-lo (0.05 ng/ml), isoalloLCA (20 μM) and TGFβ-hi (1 ng/ml) conditions with GFP^−^ naïve CD4 T cells, isolated from CD45.2 foxp3-IRES-GFP mice. **b**, Flow cytometric analysis of the FoxP3-GFP^+^ cells following *in vitro* culture. The gated cells were sorted and used for co-transfer. **c** and **d**, Rag1^-/-^ recipient mice were transferred with CD45RB^hi^ only, CD45RB^hi^ + iTreg (TGFβ-lo), CD45RB^hi^ + iTreg (isoalloLCA), or CD45RB^hi^ + iTreg (TGFβ-high) weight change monitored for eight weeks (**c**), colon length at the end of the experiment (**d**). **e-h**, Flow cytometric analysis and quantification of the percent of CD45.1 and CD45.2 (**e, f**) and the percentage of FoxP3+ cells in the CD45.2 population (**g, h**) in each condition. **i**, quantification of total CD45.1 cell number in the colon lamina propria. Error bars represent standard deviation. ns, not-significant; ^∗∗∗^p<0.001; ^∗∗^p<0.01; ^∗^p<0.05 by unpaired t-test with two-tailed p-value.

**Extended Data Figure 9.**
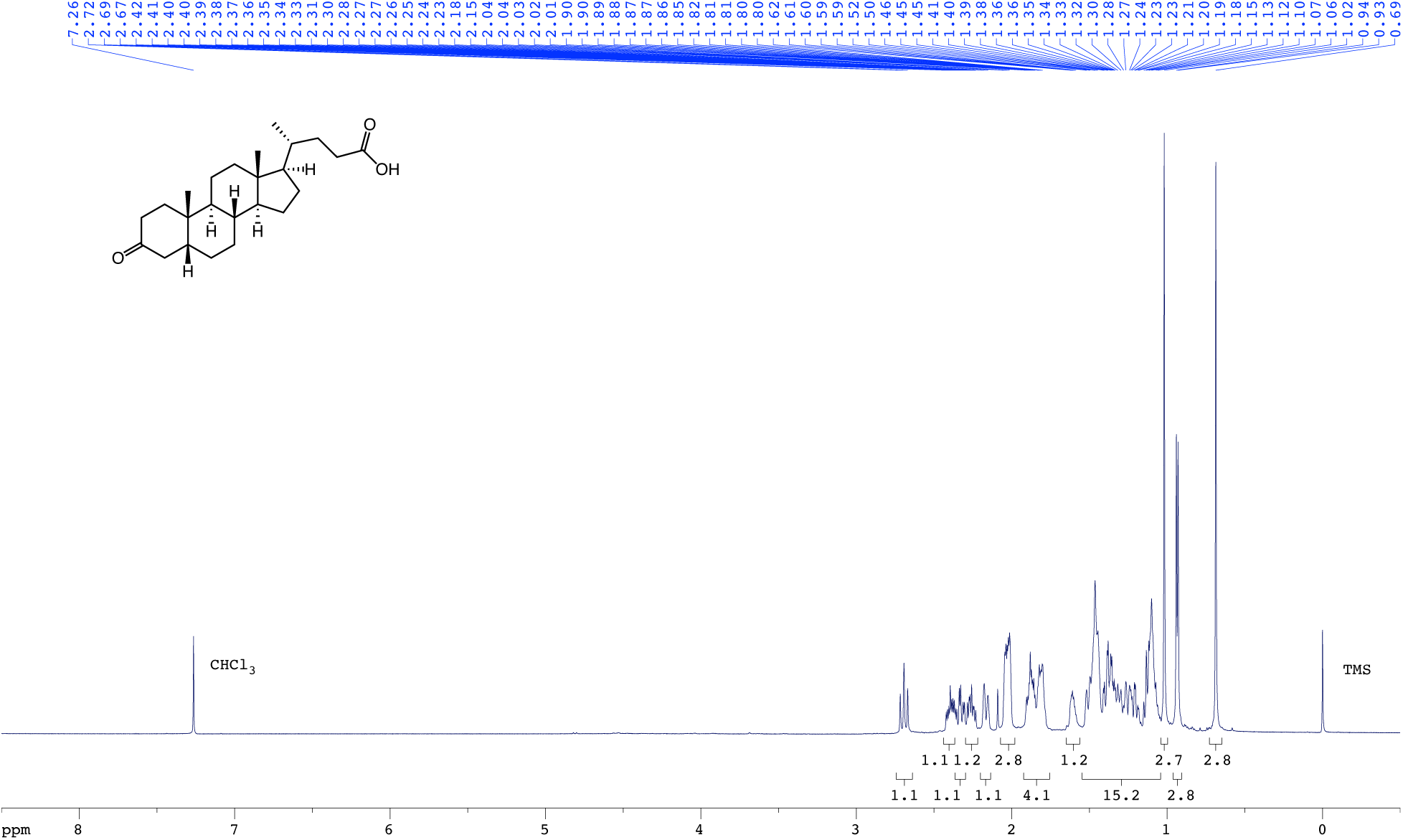
^1^H NMR spectrum (600 MHz, CDCl3) of 3-oxoLCA.

**Extended Data Figure 10.**
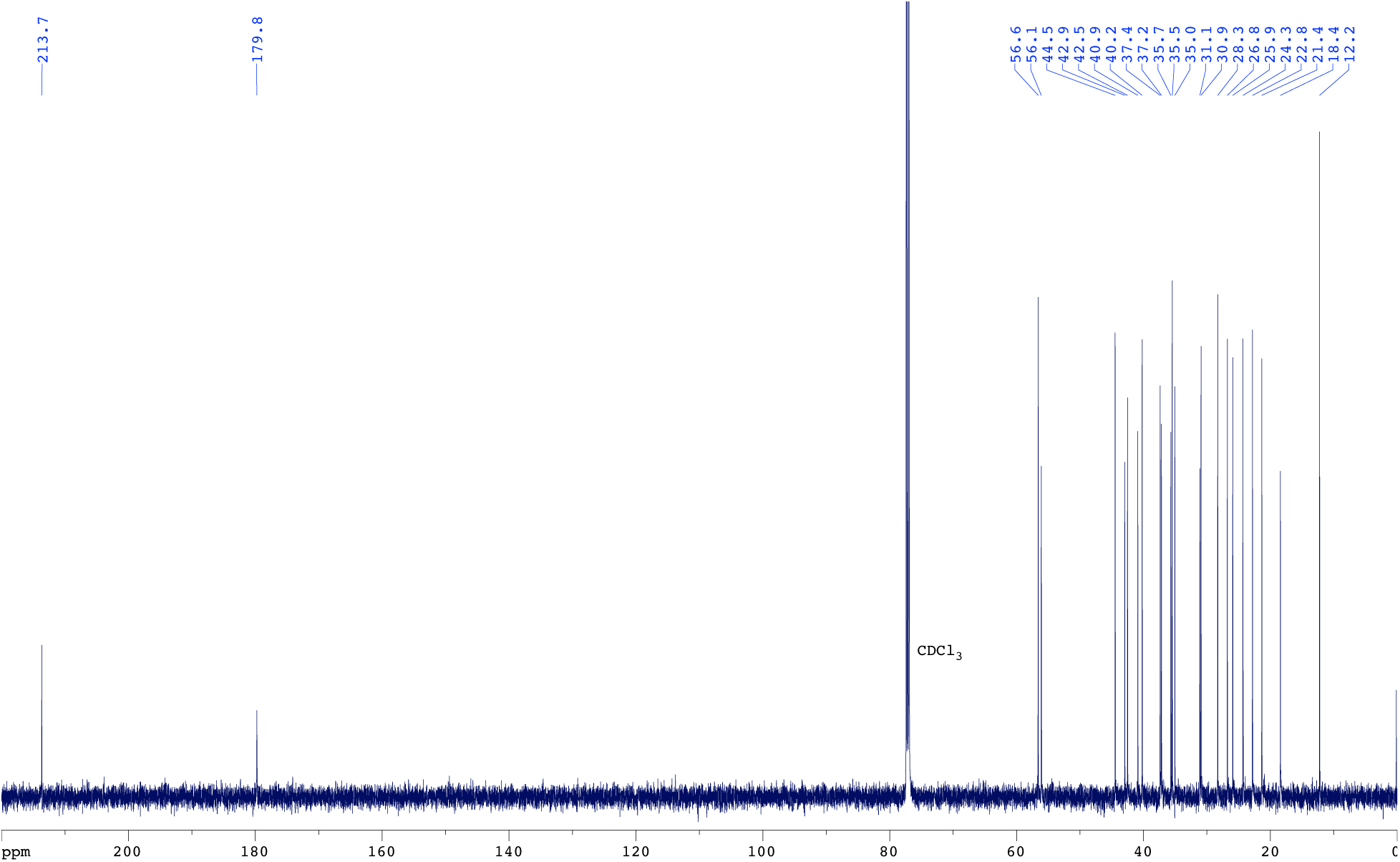
^13^C{^1^H} NMR spectrum (151 MHz, CDCl_3_) of 3-oxoLCA.

**Extended Data Figure 11.**
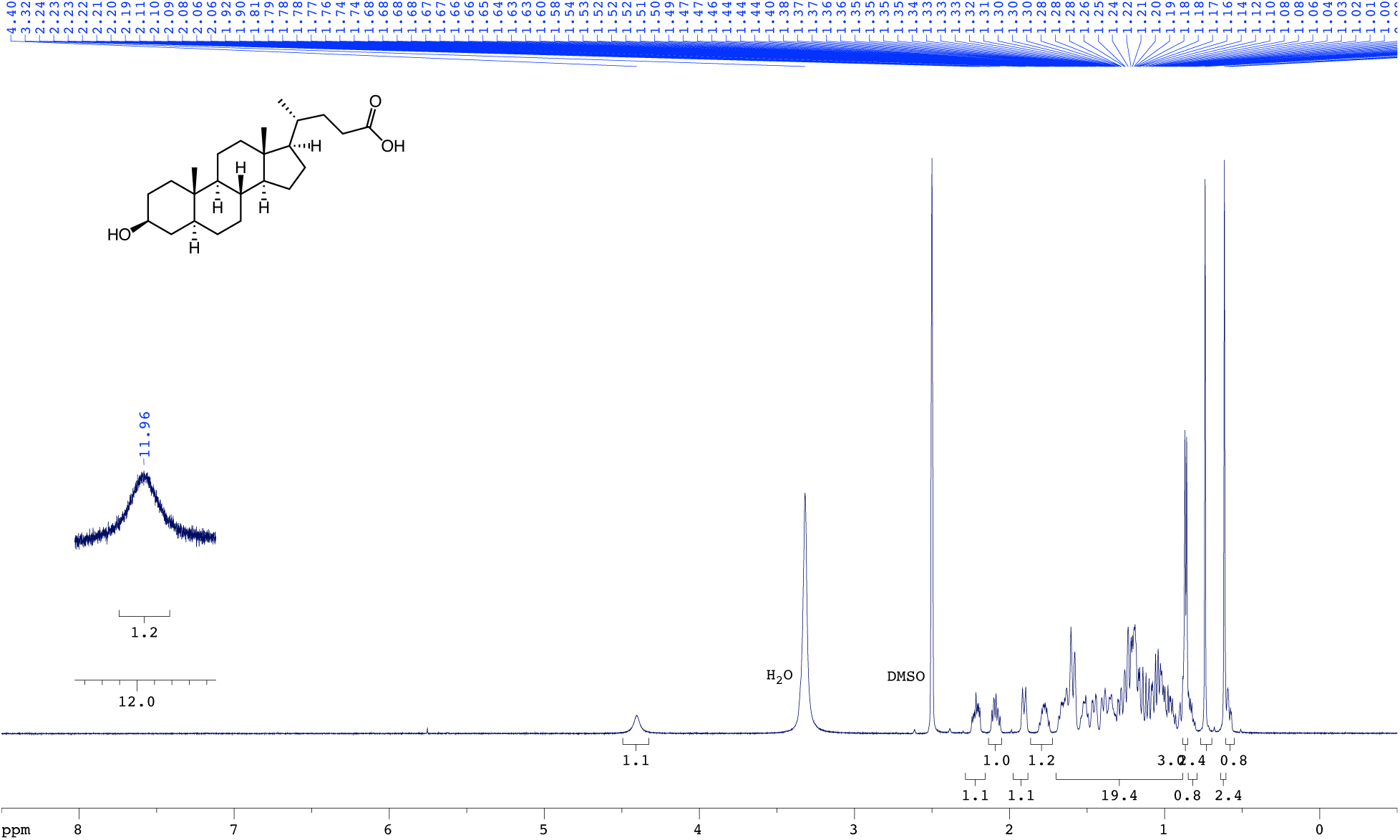
^1^H NMR spectrum (600 MHz, DMSO-*d*_6_) of isoalloLCA.

**Extended Data Figure 12.**
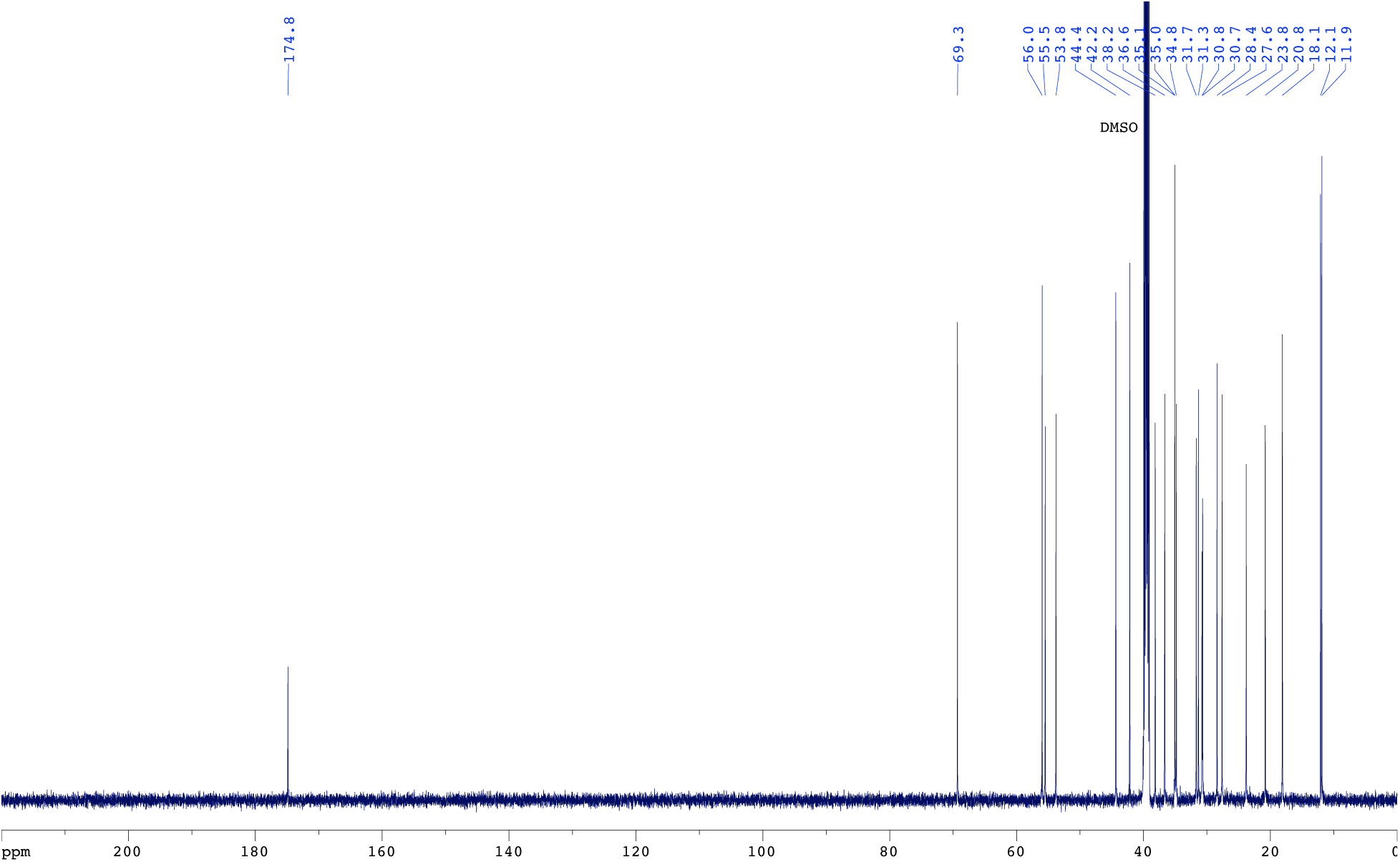
^13^C{^1^H} NMR spectrum (151 MHz, DMSO-*d*_6_) of isoalloLCA.

